# Exploration of target spaces in the human genome for protein and peptide drugs

**DOI:** 10.1101/2020.04.05.026112

**Authors:** Zhongyang Liu, Honglei Li, Zhaoyu Jin, Yang Li, Feifei Guo, Yangzhige He, Xinyue Liu, Dong Li, Fuchu He

## Abstract

**Motivation:** Protein and peptide drugs, after decades of development have grown into a major drug class of the marketplace. Target identification and validation is crucial for their discovery, and bioinformatics estimation of candidate targets based on characteristics of successful target proteins will help improve efficiency and success rate of target selection. However, owing to the development history of the pharmaceutical industry, previous systematic exploration of target space mainly focused on traditional small-molecule drugs, whereas that for protein and peptide drugs is blank. Here we systematically explored target spaces in the human genome specially for protein and peptide drugs.

**Results:** We found that compared with other proteins, targets of both successful protein and peptide drugs have their own characteristics in many aspects and are also significantly different from those of traditional small-molecule drugs. Further based on these features, we developed effective genome-wide target estimation models respectively for protein and peptide drugs.

## Introduction

Protein and peptide drugs, whose emergence has revolutionized the pharmaceutical industry, after decades of development, have grown into a major drug class of the marketplace, and are maintaining rapid development. Since the approval of the first recombinant protein drug in 1982 (Humulin, recombinant human insulin, Eli Lilly), the number of approved protein and peptide biopharmaceuticals increased rapidly, from less than 10 in the 1980s, to approval rate of 10∼12 per year which had remained constant since 1995 (by July 2014, based on United States and European Union markets) (1). Thus far, more than 200 therapeutic proteins and peptides have been approved by US Food and Drug Administration (FDA) (2). A survey from 2004 to 2014 indicated that protein and peptide biopharmaceuticals represented 21∼26% of genuinely new drug approvals (excluding biosimilars, me-too products and products previously approved elsewhere) in United States (1). Especially in 2016, FDA approved the lowest number of small-molecule drugs in nearly 45 years, resulting in that in this year the proportion of protein and peptide new molecule entity (NME) approvals reached 40%, highest to date (3). And even though 2017 and 2018 regained high total approval numbers, the high proportion of protein and peptide new drugs (37% for 2017, 30% for 2018) still remained (4,5). Meanwhile, the market value of protein and peptide drugs is rising steadily. In 2018, of the top 10 best-selling drugs, 7 were therapeuticproteinsorpeptides (https://www.genengnews.com/a-lists/top-15-best-selling-drugs-of-2018/). Outlook by EvaluatePharma (“World Preview 2017, Outlook to 2022”, 10^th^ Edition, Jun. 2017) forecasted that the market share of biologics (the vast majority of which are protein and peptide drugs) was expected to increase from 25% in 2016 to 30% in 2022, and in 2022 52% of the top 100 product sales would come from biologics.

Protein and peptide drugs, currently including hormones, growth factors, blood factors, thrombolytics and anticoagulants, interferons and interleukins, monoclonal antibodies etc., have already covered a wide range of therapeutic areas involving cancer, various inflammation-related conditions, metabolic disorders, autoimmune diseases and cardiovascular diseases etc. (1,6). The success of protein and peptide-based therapeutics over past several decades, besides the biotechnology development, stems from their several advantages. Compared with traditional small-molecule drugs, protein and peptide drugs offer excellent target specificity, high potency of action, few side-effects and low toxicity (7,8,9), and biologics have almost twice the clinical development success rate as small-molecule NMEs according to a survey from 2006 to 2015 by Biotechnology Innovation Organization (BIO) (“Clinical Development Success Rates 2006-2015”, Jun. 2016). Meanwhile, owing to their larger interaction contact surfaces, proteins and peptides are more suitable for targeting protein-protein interactions than small-molecules, which as the potential drug targets have attracted much attention in recent years (10).

Target identification and validation is crucial for protein and peptide drug discovery, and bioinformatics prediction of candidate targets (i.e. predicting whether a protein is proper to be used as a drug target) based on characteristics of successful protein/peptide drugs’ targets will be helpful for this process. An analysis of all 113 FDA-approved first-in-class drugs (which are referred to as those that modulate an - until then - unprecedented target or biological pathway) from 1999 to 2013 showed that ∼70% were identified through target-based approaches in which target identification is the first step, and especially the proportion approximated 100% for biologic drugs (11). Many experimental drug failures are attributed to inappropriate target selection, attiring a large number of resources (12,13). It is vital to provide evidence as much as possible to support a target choice before investing more resources in the target (14). Besides disease relevance, successful drug targets generally have some common features different from other non-target proteins. Systematic summary of the features of successful drug targets and further based on these features computationally estimating candidate targets will not only contribute to the understanding of cellular roles of targets and the molecular mechanism of action of drugs from an overall perspective, but also more importantly greatly increase the efficiency and success rate of target selection.

Owing to the development history of the pharmaceutical industry, up to now the work of systematical analyses of successful targets and prediction of candidate targets mainly focused on traditional small-molecule drugs, while that specially for protein and peptide drugs is still lacking. In previous studies, properties of target proteins such as protein families, biochemical functions, subcellular location, network topology, sequence etc. were explored (e.g. 14∼17). Further some target prediction methods have been developed (e.g. 14,16∼19). For example, early common prediction strategies included those based on domain family affiliation (e.g. 19) and those by searching binding pockets on the protein surface based on 3D structures to identify proteins that may bind to small-molecule drugs (e.g. 20). Recently some studies used machine learning methods on basis of summarized target features on sequence, physicochemical properties, network topology etc. to establish target prediction algorithms (e.g. 14,18,21,22). However, small-molecule drugs’ targets dominated all these studies, and systematical exploration of target space specially for protein/peptide drugs is absent.

It is necessary to systematically explore target space in the human genome specially for protein and peptide drugs, because of the potential difference between targets of different types of drugs. Realizing this potential difference, Sanseau *et al*. approximately distinguished potential targets of protein drugs from those of small-molecule drugs in genes from genome-wide association study (GWAS) using their respective features, containing signal peptide/transmembrane domain for the former and domain family distribution feature for the latter (23). Similarly, Jeon *et al*. roughly separated putative cancer targets for protein, peptide and small-molecule drugs respectively using extracellular domain inclusion, peptide-binding domain inclusion and corresponding small-molecule inhibitors existence (24). Zhu *et al*. found that the targets of biologics have higher degree and less cluster coefficient than those of small-molecule drugs (25). However, in these studies they didn’t perform any systemic analyses/comparison of targets for protein, peptide and small-molecule drugs, and didn’t construct any target prediction models.

Considering these issues, in this paper we systematically explored target spaces in the human genome specially for protein and peptide drugs (Supplementary Fig. 1). Firstly we comprehensively collected therapeutic targets of approved protein and peptide drugs. Then we systematically analyzed/compared the properties of these targets from multiple aspects to respectively reveal the features of targets of protein and peptide drugs. Finally, after feature selection by mRMR method, we used navï e Bayesian classifier to integrate several representative features to build the target prediction models separately for protein and peptide drugs.

## Materials and Methods

### Therapeutic targets

Therapeutic targets are defined as those proteins (or other biomolecules) through the interactions with which the drug exerts the therapeutic effects, excluding side-effect targets and other binding proteins without pharmacological efficacy (26,27).

We collected therapeutic targets and their corresponding approved drugs (removing withdrawn drugs) from TTD (version: 5.1.02) (28), GtoPdb (downloaded on 03/19/2017) (29), DrugBank (downloaded on 07/26/2015) (30) databases and Rask-Andersen *et al*.’s paper (26). For GtoPdb, we only considered its strictly defined “primary” targets; and for DrugBank, only drug targets with known pharmacological action were used. All targets of “group” type such as a protein complex were excluded. Drug types (protein, peptide or small-molecule drugs) were distinguished based on related annotations provided by corresponding data sources or by manual curation, where peptide drugs were separated from proteins on the basis of size, and are arbitrarily defined as molecules containing fewer than 50 amino acids (AAs) (7). Finally, 132, 55 and 634 human therapeutic target proteins uniformly represented by Swiss-Prot accession number (SP AC) respectively for protein, peptide and small-molecule drugs were obtained (Supplementary Table 1∼3).

### Golden standard datasets

To establish target prediction models, first the golden standard positive (GSP) and negative (GSN) datasets were constructed respectively for protein and peptide drugs. We take the construction of GSP and GSN sets for protein drugs as an example, and for peptide drugs, similar scheme was used (the size of GSN set was 100 for peptide drugs). The GSP set for protein drugs was directly composed of the therapeutic targets of approved protein drugs collected above. For the GSN set, considering the difficulty gaining an experimentally negative dataset, we adopted the following construction scheme. First we removed known protein drugs’ targets as many as possible from the whole human genome. To ensure the “purity” of the GSN set as “high” as possible, here the removed known targets not only included the GSP set (i.e. the therapeutic targets of approved protein drugs collected above), but also other possible targets. These other possible targets included targets of trial protein drugs from Rask-Andersen *et al*.’s paper (26), non-”primary” targets of approved protein drugs in GtoP<u>db</u> (“primary” ones had been included in the GSP set) (downloaded on 03/19/2017), and all targets (also including those without known pharmacological action) of all protein drugs (also including experimental drugs) in DrugBank (downloaded on 07/26/2015), as well as those of “group” type such as targets of protein complex type. Then from the remaining ones we randomly picked 200 proteins to construct the GSN set. Further considering the randomness during the construction of the GSN set, to avoid potential bias, we repeated the construction process of the GSN set 100 times, and finally reported mean ± standard deviation (SD) of results of 100 times. The ROC curve of 10-fold cross-validation in this paper was drawn based on the GSP set and a random GSN set.

### Independent test datasets

To assess the performance of prediction models, besides the cross-validation, three independent test sets were also adopted.

**The independent test set1:** The first independent test positive set was composed of the newly added therapeutic target set of approved protein (/peptide) drugs from DrugBank of the latest version (Release on 2018-07-03), and after removing the overlap with the GSP and GSN set, 17 (/16) targets were remained to constitute this independent positive test set for protein (/peptide) drugs. **The independent test set2:** The second independent positive test set from Rask-Andersen *et al*.’s paper (26) was composed of 125 (/51) targets of clinical trial protein (/peptide) drugs (after removing the overlap with the GSP and GSN set). **The independent test set3:** In the third independent test scheme, for protein/peptide drugs’ targets, we divided the golden standard positive set into two parts. One part composed of targets of drugs with approval time before 2010 (as well as those without approval time information) together with the GSN built above was used to train the prediction model, and the other involving new therapeutic targets introduced by drugs approved in or after 2010 was used as the independent positive test set. The approval time information of drugs in GSP sets was from related annotations of corresponding data sources or identified by Drugs@FDA (https://www.accessdata.fda.gov/scripts/cder/daf/).

The independent negative test set for each of these three groups of positive test sets above was respectively established in a similar way to the construction of the GSN set. Especially, before randomly choosing proteins of similar number as that of the corresponding independent positive test set from the human genome, besides removing the known protein/peptide drug targets, the corresponding GSN set and the independent positive test set were also excluded. Considering the randomness of the independent negative test set, we also repeated its construction process 100 times, and finally the assessment results of prediction models on independent test sets were mean ± SD of 10000 results (i.e. those of 100 prediction models constructed based on 100 golden standard datasets, on 100 independent test sets). The ROC curve based on the independent test set was drawn based on a random GSN set and a random independent test negative set.

### Data sources and methods related to property analyses of therapeutic targets

Amino acids were classified into 9 groups, including tiny (A+C+G+S+T), small (A+B+C+D+G+N+P+S+T+V),aliphatic(A+I+L+V),aromatic(F+H+W+Y), non-polar (A+C+F+G+I+L+M+P+V+W+Y), polar (D+E+H+K+N+Q+R+S+T+Z), charged (B+D+E+H+K+R+Z), basic (H+K+R) and acidic (B+D+E+Z). We used Pepstats program of EMBOSS (version 6.5.0) to count the percentage of each class of AA and the charge of a protein sequence (31). The grand average of hydropathy (GRAVY) of a protein is computed as the sum of hydropathy values (32) of its all AAs, divided by its AA sequence length. The GRAVY value and theoretical isoelectric point (pI) were calculated by ProtParam program downloaded from Comprehensive Perl Archive Network (CPAN) (https://www.perl.org/cpan.html). Amino acid (AA) sequences of human proteins were from SwissProt (downloaded on 01/11/2015) (33).

Pfam domain (34) assignments of human proteins were parsed from SwissProt (downloaded on 03/23/2016). Intrinsically disordered proteins are those that lack fixed or ordered 3-D structures. The disorder score of a protein is computed as the ratio of the length of disordered regions to its total length (35). FoldIndex was used to predict the intrinsic disorder of a protein (using default parameters) (36), and to reduce the false positive rate only those disordered regions with length no smaller than 30 were considered (35). A PEST region is a peptide sequence enriched in proline (P), glutamic acid (E), serine (S) and threonine (T), and is invariably found in proteins with a short half-life and thus hypothesized to serve as proteolytic signals (37). Here we adopted EMBOSS epestfind program (using default parameters) (31) to count the number of PEST motifs in a protein, and only “potential” PEST motifs were included. A signal peptide is a short peptide present at the N-terminus of proteins that are targeted to the endoplasmic reticulum and eventually destined to be either secreted, extracellular or periplasmic etc. (33). Signal peptides and transmembrane regions of human proteins were both parsed from related annotations of SwissProt (downloaded on 02/11/2016).

The tissue specificity score (TSPS) was adopted to measure the degree of tissue specific expression of a gene (38). See Supplementary Methods for its formula. The larger TSPS is, the more tissue-specific the expression of the gene is. We used RNA-sequencing data from 32 tissues provided by Uhlén *et al*. to compute the TSPS (39).

The data on evolutionary rates and original ages of human proteins were from our previous work (40). Human proteins were grouped into 16 age classes, from the oldest “Cellular organisms” class of age 16 composed of proteins that originated from the common ancestor of three domains of tree of life (Eukiaryota, Bacteria and Archaea) to the youngest “Homo sapiens” class of age 1 whose proteins were only found in human. We used *C*_*ratio*_ to check the gene polymorphism, which is computed as 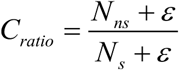 where *N*_*ns*_ and *N*_*s*_ are respectively the number of nonsynonymous and synonymous single-nucleotide polymorphisms (SNPs) in the gene and *ε* was set to 0.01 as Yao *et al*. did in order to eliminate statistical aberrations induced by relatively small sample size (21). Coding-region SNP data were from dbSNP database (downloaded on 10/18/2016) (41) and only SNPs with global minor allele frequency (GMAF) no smaller than 1% were considered.

The list of human transcriptional factors (TFs) was from Lambert *et al*.’s paper (42), involving 1639 genes. Housekeeping genes are those detected in all tissues, which were obtained based on RNA-sequencing data in 32 tissues from Uhlén *et al*.’s paper (39), involving 8874 genes. The two lists of signaling molecules and self-interacting proteins were respectively from the signal transduction network and the integrated PPI network described in the next paragraph. The human gene list coding enzymes were parsed from SwissProt (downloaded on 02/11/2016), G protein-coupled receptors (GPCRs) and kinases from UniProt (http://www.uniprot.org/docs/7tmrlist and http://www.uniprot.org/docs/pkinfam) (Release: 2017_10 of 25-Oct-2017) (33), both ion channels and nuclear hormone receptors (NHR) from HUGO Gene Nomenclature Committee (HGNC) database (downloaded on Jul. 2017) (43), and transporters from Human Transporter Database (HTD) (version: 2014-01-01) (44). Human gene-reaction associations were from Recon 2 (version: 11.05.2015) (45), and biological pathways from KEGG (downloaded on 7/13/2016) (46). In a network, degree reflects a node’s local importance, while betweenness centrality captures the degree to which the node influences the communication between other nodes in the network (47) (see Supplementary Methods for their formulas). The integrated human PPI network was obtained by integrating experimental PPIs of “direct interaction” type from Database of Interacting Proteins (DIP) (version: 4/30/2016) (48), Molecular INTeraction database (MINT) (downloaded on 7/17/2016) (49), IntAct (version: 2016-07-06) (50) and Biological General Repository for Interaction Datasets (BioGRID) (version: 3.4.138) (51), containing 69357 PPIs between 12558 proteins. The human signal transduction network was provided by Cui *et al*. (version 6) (52) and the transcriptional regulation network from Chouvardas *et al*.’s paper (53). Here the PPI and signal transduction networks were regarded as undirected networks, while the transcriptional regulation network a directed network. In the directed transcriptional regulation network, indegree (of a target gene) denotes the number of TFs regulating this target gene while outdegree (of a TF) the number of target genes regulated by this TF.

Domain enrichment ratio (DER) was used to measure the enrichment degree of a domain in the GSP set. DER is calculated as the ratio of probability of observing this domain in the GSP set to that in the whole genome. To avoid “self-circulation” problem, in this paper 10-fold cross-validation scheme was adopted to compute the Likelihood Ratio (LR) of feature DER. Nine tenths of the GSP set was used to define the enriched domains and compute corresponding DER, and the remaining one was used for test. This process was repeated ten times and the ten test results were combined to compute LR.

### Methods related to prediction model construction and estimation

We used minimum redundancy maximum relevance (mRMR) feature selection method for features selection. mRMR ranks features based on both their relevance to the classification variable and the redundancy between each other (47,54) (see Supplementary methods for details). After feature selection, navï e Bayes classifier was used to integrate multiple features to establish the target prediction model (see Supplementary methods for details). Finally we used Receiver Operating Characteristic (ROC) curve to measure the performance of the prediction model, based on 10-fold cross-validation and independent test sets (see Supplementary methods for details).

## Results

### The collection of known therapeutic targets

The main aim of our work is to summarize the features of successful drug targets and further based on these features to provide quantitative advice for the future target selection, and thus the collection scope of known drug targets is crucial. Here we only collected therapeutic targets of approved drugs (see Materials and Methods). In total we obtained 132, 55 and 634 human therapeutic target proteins respectively for approved protein, peptide and small-molecule drugs (Supplementary Table 1∼3).

Firstly, we noticed that although there was overlap between therapeutic targets of protein, peptide and small-molecule drugs (that is, there were some proteins that can be used as targets of two or three types of drugs), most targets were specific to respective kind of drugs (Fig. 1a), suggesting potential difference between target spaces of different types of drugs.

**Figure 1.**
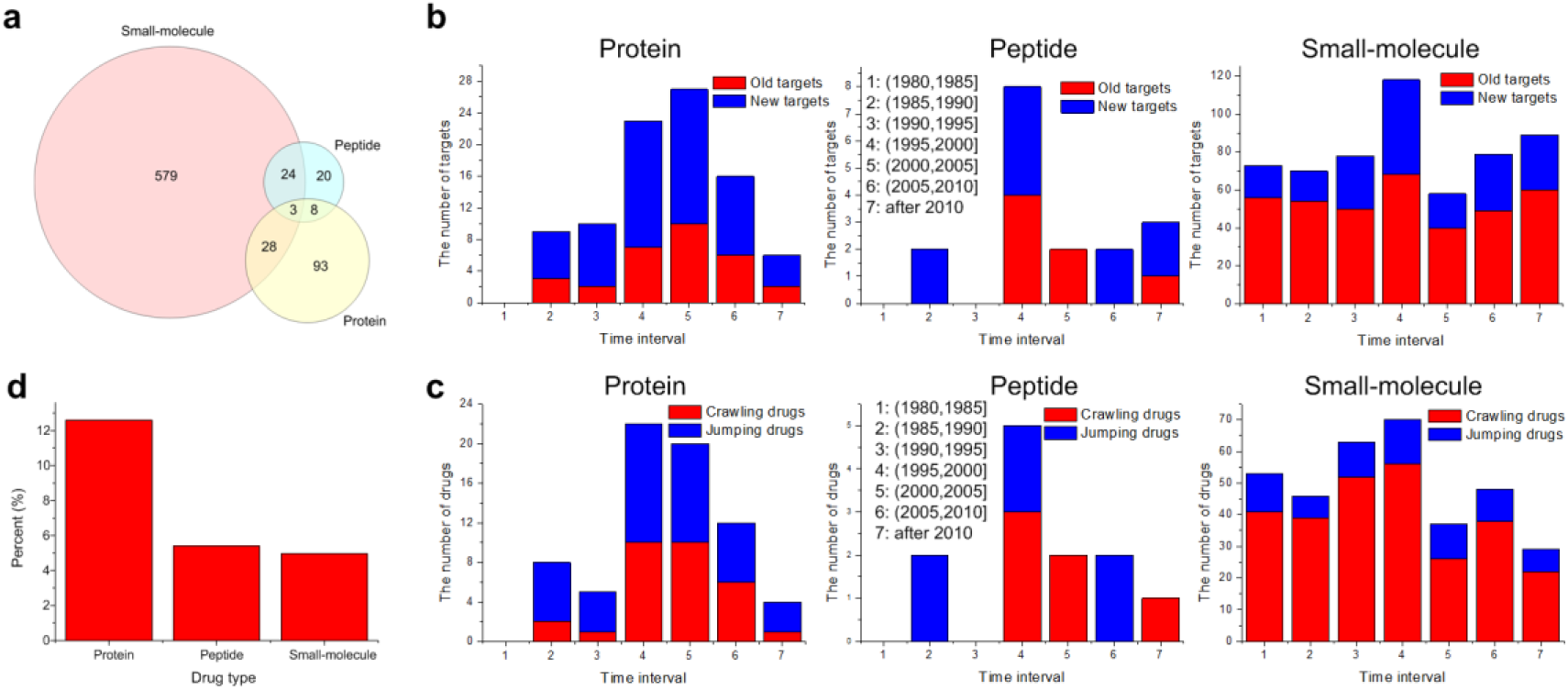
Statistics of therapeutic targets of approved protein, peptide and small-molecule drugs. **a** The Venn graph of therapeutic targets of the three types of drugs. The number of each part on the graph is the number of targets. **b** The composition of therapeutic targets of new drugs approved each five years. “Old targets” are referred to as those that have been used as targets of previously approved drugs, while “new targets” not. Each five years is a bin, and the year scope of each bin is described on the graph. **c** The composition of drugs newly approved each five years. We used Yildirim *et al*.’s definition (55), and “jumping drugs” are referred to as those with totally new target sets, whereas “crawling drugs” with at least one “old target”. The analyses of **b** and **c** only included drugs with at least one therapeutic target, and these results were obtained based on approved drugs’ therapeutic target data from Rask-Andersen *et al*.’s paper (26) (see Materials and Methods). We obtained consistent conclusion based on data from DrugBank (downloaded on 07/26/2015, see Materials and Methods) (Supplementary Fig. 2). **d** The percent of known drug-disease associations satisfying the condition that at least one therapeutic target of the drug is simultaneously its corresponding disease’s known related gene. Only known drug-disease associations containing drugs with at least one target and diseases with at least one known related gene were involved in this analysis (see Supplementary Materials and Methods).

Next, we added approval time labels for drugs, and from target perspective compared the innovation power of three different types of drugs. In 2007 Yildirim *et al*. confirmed that the pharmaceutical industry tends to target already validated target proteins based on drug-target interaction data which merged protein, peptide and small-molecule drugs (55). Here we found that for protein drugs, averagely over half of (68.06%) targets of drugs newly approved every five years were “new targets” which have not been used as targets before by any approved drug, while the average proportion (32.29%) was much smaller for small-molecule drugs (rank sum test, P-value=0.0010) (Fig. 1b). The analyses of the new drug composition also led to consistent tendency, that is, the average percent of “jumping drugs” was much higher for protein drugs than that for small-molecule drugs (64.09%>21.43%, rank sum test, P-value=0.0010) (Fig. 1c), where those new drugs whose all targets are “new targets” are called as “jumping drugs”, otherwise “crawling drugs” (55). As for the peptide drugs, although the average proportion of “new targets” (Fig. 1b) or “jumping drugs” (Fig. 1c) was larger than that of small-molecule drugs, we did not obtain statistical significance. These results indicate that protein new drugs tend to bind “new targets” rather than validated targets, and have stronger target innovation power than traditional small-molecule drugs. And Yildirim *et al*.’s conclusion that new drugs tend to target already validated target proteins mentioned above (55) could be attributed to dominant small-molecule drugs.

At last, we analyzed and compared the drug-disease relationship for different types of drugs by simply counting the proportion of known drug-disease associations satisfying the condition that at least one of the drug’s therapeutic targets is also the known related gene of the corresponding disease (see Supplementary Methods). We found that the proportion for protein drugs was significantly larger than that for small-molecule drugs (12.60%>4.96%, Fisher’s exact test, P-value<10^−4^) (Fig. 1d). This indicates that compared with small-molecule drugs, protein drugs tend to directly target disease-causing proteins, which confirms the previous opinion to some degree that compared with small-molecule drugs, protein drugs tend to cure the diseases rather than only treat their symptoms (9). But here we did not see the significant difference between peptide and small-molecule drugs (Fig. 1d).

### Distinctive properties of targets of protein and peptide drugs

Based on the obtained lists of successful therapeutic targets, in this part we intend to answer two questions: 1) what characteristics protein and peptide drugs’ targets respectively have, compared with other proteins; 2) whether and how they are different from conventional small-molecule drugs’ targets. We checked multiple aspects of properties to give the answer.

For the first question, we found that successful protein drugs’ targets, compared with other non-target proteins, in the sequence and physicochemical property aspect, tend to contain significantly higher proportion of small and aromatic amino acids (AAs), lower basic and charged AAs, less polar and more non-polar AAs and thus significantly more hydrophobic (measured by GRAVY, see Materials and Methods), and their charges tend to be negative and therefore have significantly lower theoretical pI (Table 1).

**Table 1.**
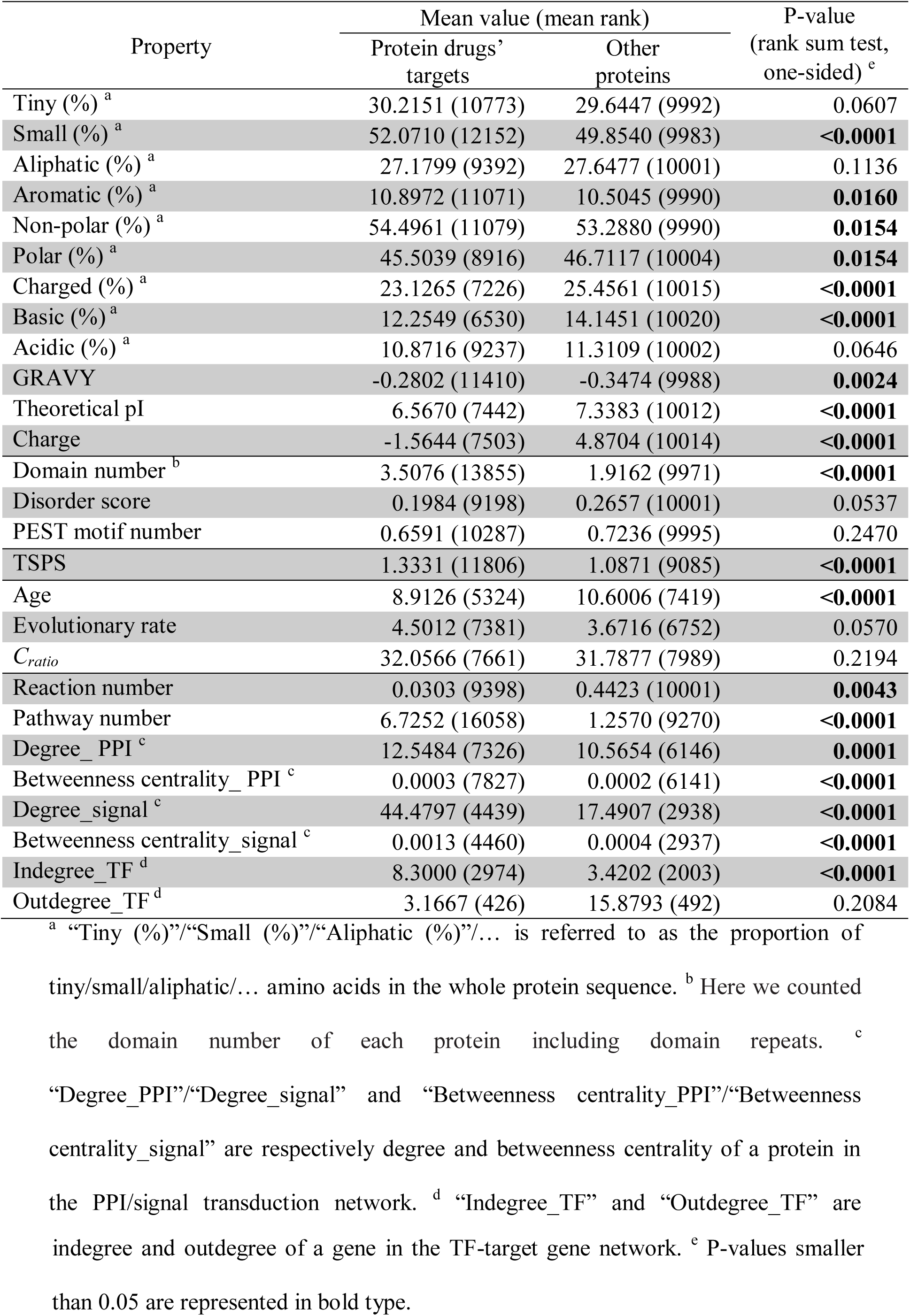
The difference of quantitative properties between protein drugs’ targets and other proteins

| Property | Mean value (mean rank) |  | P-value<br>(rank sum test,<br>one-sided) <sup>e</sup> |
| --- | --- | --- | --- |
|  | Protein drugs' targets | Other proteins |  |
| Tiny (%) <sup>a</sup> | 30.2151 (10773) | 29.6447 (9992) | 0.0607 |
| Small (%) <sup>a</sup> | 52.0710 (12152) | 49.8540 (9983) | **<0.0001** |
| Aliphatic (%) <sup>a</sup> | 27.1799 (9392) | 27.6477 (10001) | 0.1136 |
| Aromatic (%) <sup>a</sup> | 10.8972 (11071) | 10.5045 (9990) | **0.0160** |
| Non-polar (%) <sup>a</sup> | 54.4961 (11079) | 53.2880 (9990) | **0.0154** |
| Polar (%) <sup>a</sup> | 45.5039 (8916) | 46.7117 (10004) | **0.0154** |
| Charged (%) <sup>a</sup> | 23.1265 (7226) | 25.4561 (10015) | **<0.0001** |
| Basic (%) <sup>a</sup> | 12.2549 (6530) | 14.1451 (10020) | **<0.0001** |
| Acidic (%) <sup>a</sup> | 10.8716 (9237) | 11.3109 (10002) | 0.0646 |
| GRAVY | -0.2802 (11410) | -0.3474 (9988) | **0.0024** |
| Theoretical pI | 6.5670 (7442) | 7.3383 (10012) | **<0.0001** |
| Charge | -1.5644 (7503) | 4.8704 (10014) | **<0.0001** |
| Domain number <sup>b</sup> | 3.5076 (13855) | 1.9162 (9971) | **<0.0001** |
| Disorder score | 0.1984 (9198) | 0.2657 (10001) | 0.0537 |
| PEST motif number | 0.6591 (10287) | 0.7236 (9995) | 0.2470 |
| TSPS | 1.3331 (11806) | 1.0871 (9085) | **<0.0001** |
| Age | 8.9126 (5324) | 10.6006 (7419) | **<0.0001** |
| Evolutionary rate | 4.5012 (7381) | 3.6716 (6752) | 0.0570 |
| *C<sub>ratio</sub>* | 32.0566 (7661) | 31.7877 (7989) | 0.2194 |
| Reaction number | 0.0303 (9398) | 0.4423 (10001) | **0.0043** |
| Pathway number | 6.7252 (16058) | 1.2570 (9270) | **<0.0001** |
| Degree_PPI <sup>c</sup> | 12.5484 (7326) | 10.5654 (6146) | **0.0001** |
| Betweenness centrality_PPI <sup>c</sup> | 0.0003 (7827) | 0.0002 (6141) | **<0.0001** |
| Degree_signal <sup>c</sup> | 44.4797 (4439) | 17.4907 (2938) | **<0.0001** |
| Betweenness centrality_signal <sup>c</sup> | 0.0013 (4460) | 0.0004 (2937) | **<0.0001** |
| Indegree_TF <sup>d</sup> | 8.3000 (2974) | 3.4202 (2003) | **<0.0001** |
| Outdegree_TF <sup>d</sup> | 3.1667 (426) | 15.8793 (492) | 0.2084 |

<sup>a</sup> “Tiny (%)”/“Small (%)”/“Aliphatic (%)”/... is referred to as the proportion of tiny/small/aliphatic/... amino acids in the whole protein sequence. <sup>b</sup> Here we counted the domain number of each protein including domain repeats. <sup>c</sup> “Degree\_PPI”/“Degree\_signal” and “Betweenness centrality\_PPI”/“Betweenness centrality\_signal” are respectively degree and betweenness centrality of a protein in the PPI/signal transduction network. <sup>d</sup> “Indegree\_TF” and “Outdegree\_TF” are
indegree and outdegree of a gene in the TF-target gene network. <sup>c</sup> P-values smaller than 0.05 are represented in bold type.

In the structural aspect, compared with other proteins, protein drugs’ targets contain significantly more domains (Table 1), indicating they tend to have more complex structure and functions. Further we observed that compared with other proteins, significantly more protein drugs’ targets contain the transmembrane region and signal peptide (Table 2), which is consistent with the previous knowledge (23).

**Table 2.**
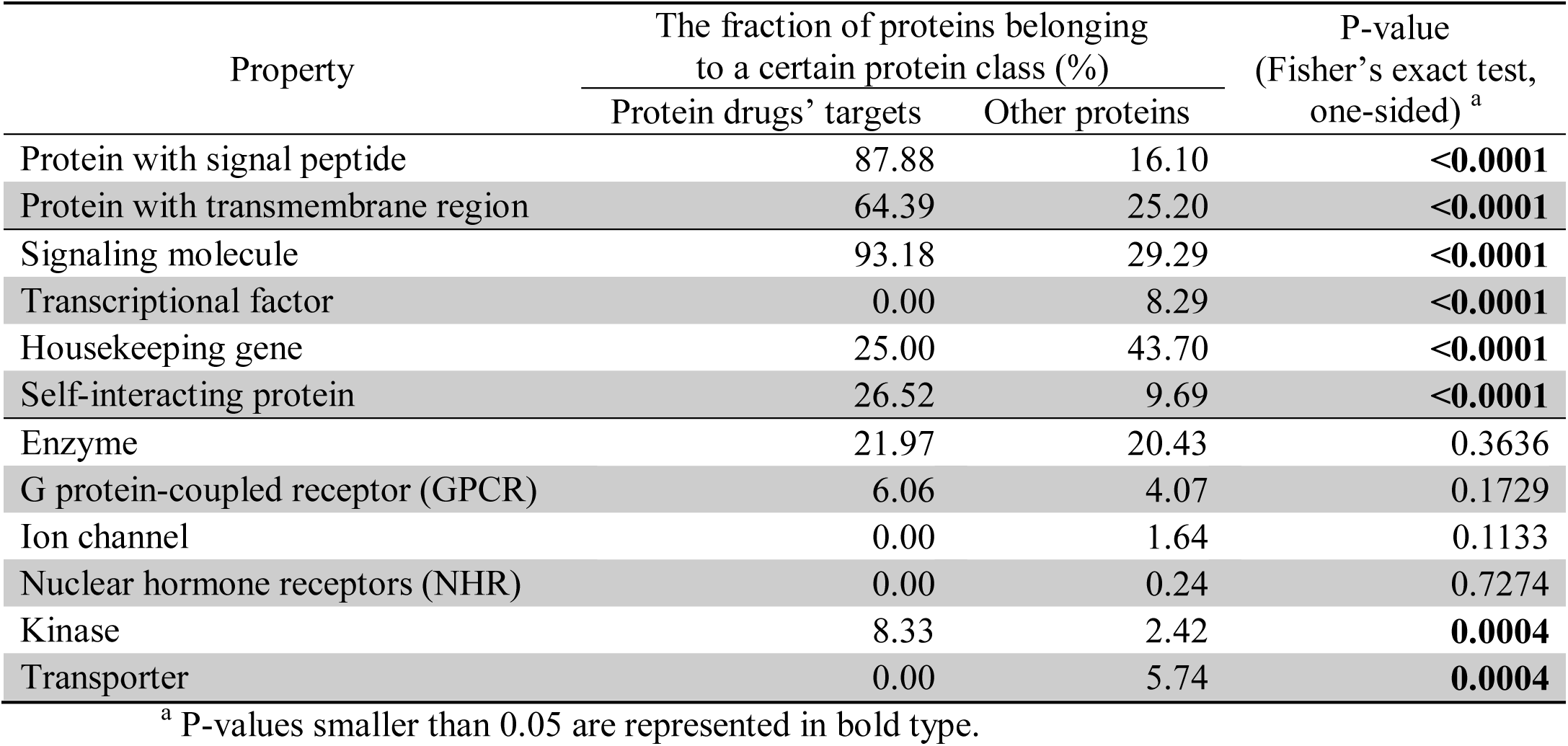
The difference of qualitative properties between protein drugs’ targets and other proteins

| Property | The fraction of proteins belonging to a certain protein class (%) |  | P-value<br>(Fisher's exact test, one-sided) <sup>a</sup> |
| --- | --- | --- | --- |
|  | Protein drugs' targets | Other proteins |  |
| Protein with signal peptide | 87.88 | 16.10 | **<0.0001** |
| Protein with transmembrane region | 64.39 | 25.20 | **<0.0001** |
| Signaling molecule | 93.18 | 29.29 | **<0.0001** |
| Transcriptional factor | 0.00 | 8.29 | **<0.0001** |
| Housekeeping gene | 25.00 | 43.70 | **<0.0001** |
| Self-interacting protein | 26.52 | 9.69 | **<0.0001** |
| Enzyme | 21.97 | 20.43 | 0.3636 |
| G protein-coupled receptor (GPCR) | 6.06 | 4.07 | 0.1729 |
| Ion channel | 0.00 | 1.64 | 0.1133 |
| Nuclear hormone receptors (NHR) | 0.00 | 0.24 | 0.7274 |
| Kinase | 8.33 | 2.42 | **0.0004** |
| Transporter | 0.00 | 5.74 | **0.0004** |

<sup>a</sup> P-values smaller than 0.05 are represented in bold type.

In the gene expression and evolutionary aspect, protein drugs’ targets tend to be tissue-specific (which is measured by TSPS, see Materials and Methods) (Table 1), which may avoid more potential side-effects. Further protein drugs’ targets originated significantly later. And although they evolve more quickly, the difference does not reach statistical significance (P-value=0.0570, Table 1). There is also no apparent difference between protein drugs’ targets and other proteins in gene nonsynonymous polymorphism (measured by *C*_*ratio*_, see Materials and Methods) at the human population level (Table 1).

Furthermore in the functional aspect, 93% of the collected protein drugs’ targets participate in signal transduction, several times higher than that of other proteins, but transcriptional factors are apparently depleted in them (Table 2). Housekeeping genes are also significantly lacked (Table 2), consistent with the high tissue specificity of protein drugs’ targets. And protein drugs’ targets are significantly enriched with self-interacting proteins (Table 2), in line with our previous observation (47). In biochemical function aspect, protein drugs’ targets are of special characteristics. Previous studies summarized that the vast majority of known drug targets, which are mainly composed of small-molecule drugs’ targets, belong to biochemical groups including enzyme, GPCR, ion channel, transporter, kinase and NHR (e.g. 19,29). However here we saw that unlike traditional small-molecule drugs’ targets, protein drugs’ targets are not enriched with enzymes, GPCRs, ion channels and NHRs, and even significantly lack transporters (Table 2), suggesting their potential great difference from small-molecule drugs’ targets.

At last in pathway and network analysis aspect, compared with other proteins, first we observed that protein drugs’ targets are involved in significantly less metabolic reactions (Table 1), which indicates again that unlike small-molecule drugs, current protein drugs do not tend to intervene metabolic processes. Further protein drugs’ targets tend to participate in more biological pathways, topologically occupy more important positions in the protein-protein interaction (PPI) network and the signal transduction network (Table 1). At last in the transcriptional regulation network we found that protein drugs’ targets are regulated by significantly more TFs (Table 1), which is consistent with their tissue specificity since a high level of transcriptional regulation may be needed for tissue-specific genes (56).

For the second question, by comparison we saw that indeed in many aspects successful protein drugs’ targets are significantly different from those of conventional small-molecule drugs including AA composition and physicochemical properties, protein structure, evolution, network topology and especially functions (Supplementary Table 4 and 5). For example, protein drugs’ targets contain significantly lower proportion of non-polar and higher polar AAs and thus are more hydrophilic than small-molecule drugs’ targets, and have apparently higher structural disorder (Supplementary Table 4). In evolution, protein drugs’ targets tend to be significantly younger and evolve faster, and more nonsynonymously polymorphic in human population (Supplementary Table 4). In the transcriptional regulation network, compared with small-molecule drugs’ targets, protein drugs’ targets tend to be regulated by significantly more TFs but regulate significantly less other genes as TFs (Supplementary Table 4), which might partly explain the fewer side-effects of protein drugs. Especially in the aspect of function, two types of drug targets have significantly different biochemical distribution, and protein drugs’ targets have significantly lower proportion of enzymes, GPRCs, ion channels, NHRs and transporters, compared with small-molecule drugs’ targets (Supplementary Table 5). As for peptide drugs’ targets, compared with either other non-target proteins or small-molecule drugs’ targets, we found that they also have distinctive characteristics in many aspects (Supplementary Table 6 and 7; Supplementary Table 8 and 9). And meanwhile peptide and protein drugs’ targets are also distinguishable on many properties (Supplementary Table 10 and 11).

In summary, both successful protein and peptide drugs’ targets have their own characteristics. Meanwhile targets of protein, peptide and small-molecule drugs are significantly different between each other in many aspects. Therefore it is quite necessary to respectively construct target prediction models specially for currently highly-concerned protein and peptide drugs, and meanwhile the establishment of efficient prediction models is also feasible based on their these distinctive characteristics.

### Effective prediction of targets of protein and peptide drugs

To build prediction models, efficient prediction features were first collected. Theoretically, all the features of protein/peptide drugs’ targets with statistical significance revealed above can be efficiently used for prediction, and this was confirmed by Likelihood Ratio (LR) results (see Supplementary Methods) (Supplementary Fig. 3 and 4). Besides, considering that successful protein/peptide drugs’ targets may tend to contain special domains, we also predicted protein/peptide drugs’ targets by domain enrichment ration (DER) (see Materials and Methods). A strong correlation between LR and DER indicated that as we expected this feature is also a qualified predictor (Supplementary Fig. 3ab and Supplementary Fig. 4w).

These collected features were confirmed to be efficient, however apparently their prediction abilities are different (Supplementary Table 12) and more importantly there is some redundancy between each other (Supplementary Table 13). Therefore feature selection is necessary. We used mRMR method to rank these features, based on both their efficiency and redundancy between each other (see Supplementary Methods) (Supplementary Table 14 for protein drugs and Supplementary Table 15 for peptide drugs).

Finally we adopted navï e Bayes rule to integrate these features to construct the target prediction model for protein/peptide drugs. Taking the model construction for protein drugs as an example, starting from the most powerful feature “DER”, according to the feature order ranked by mRMR (Supplementary Table 14), we added features into the integrated prediction model one by one. Ultimately 28 models were obtained, from Model_1 only composed of “DER” to Model_28 integrating all features (Fig. 2). From 10-fold cross-validation results we observed that as we expected integrating more features into the model did not achieve stronger prediction efficacy. By and large, the AUC was gradually improved as the features were added one by one, but we saw that after the integrated features’ number exceeded eight, the performance kept unchanged or even declined (Fig. 2). It is confirmed again that good classification performance can be obtained only by integrating several “representative” features with relatively high efficacy and low redundancy (47). Finally we used the “Model_8_protein”, integrating eight features, including DER, “transcriptional factor”, “signaling molecule”, “signal peptide”, “transmembrane region”, “pathway number”, “indegree_TF” and “betweenness centrality_PPI”, as our prediction model of protein drugs’ targets, which achieved an outstanding AUC of 0.9654 ± 0.0047. In the similar way, we established the model for peptide drugs - “Model_4_peptide” - integrating DER, “signal peptide”, “transmembrane region” and “signaling molecule”, whose AUC reached 0.9370 ± 0.0150, also displaying its excellent prediction ability (Supplementary Fig. 5).

**Figure 2.**
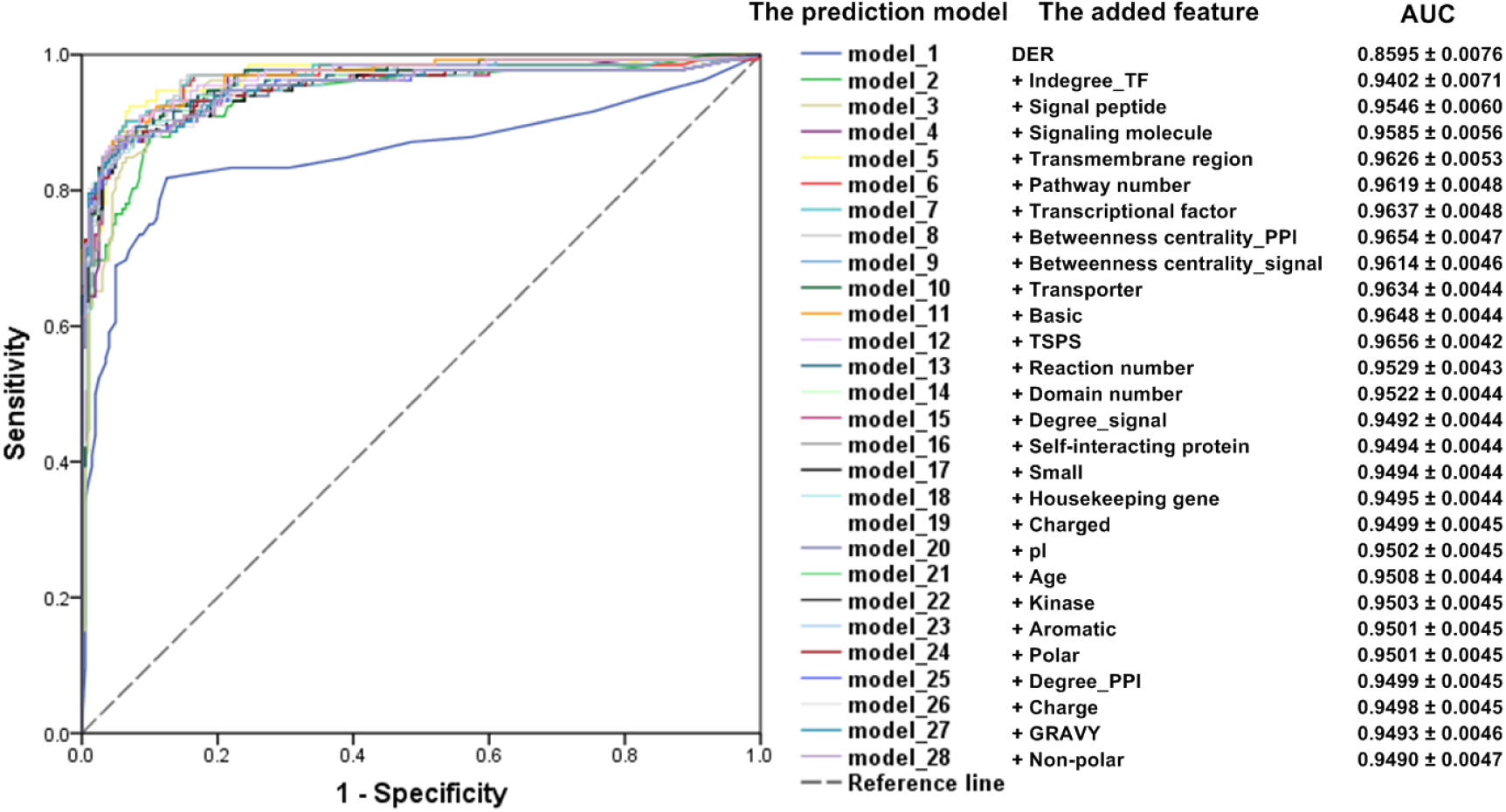
ROC curves and their corresponding AUCs of target prediction models for protein drugs integrating *n* features based on 10-fold cross-validation. ROC curves of these models are plotted in different colors.

Besides 10-fold cross-validation, three independent test datasets were also used to evaluate the performance of prediction models. As seen in Fig. 3, the constructed “Model_8_protein” and “Model_4_peptide” can well discriminate therapeutic targets of approved protein/peptide drugs newly added by DrugBank of the latest version (Release on 2018-07-03) (Fig. 3a and 3b) and clinical trial drug targets collected by Rask-Andersen *et al*. (26) (Fig. 3c and 3d) from other non-target proteins (see Materials and Methods). Especially in the third test, the GSP set was divided into two subsets. One was composed of therapeutic targets of drugs with approval time before 2010, and the other included new targets brought by drugs approved in or after 2010. The former was used to train “Model_8_protein”/”Model_4_peptide”, and the latter as the test set. This scheme was designed to stimulate the real development process of drug targets, in order to approximately estimate the performance of our prediction system on future truly successful targets. In result”Model_8_protein” and “Model_4_peptide” trained based on this scheme still obtained satisfactory performance (Fig. 3e and 3f), strongly showing effectiveness of our prediction system on distinguishing true targets from other non-target proteins.

**Figure 3.**
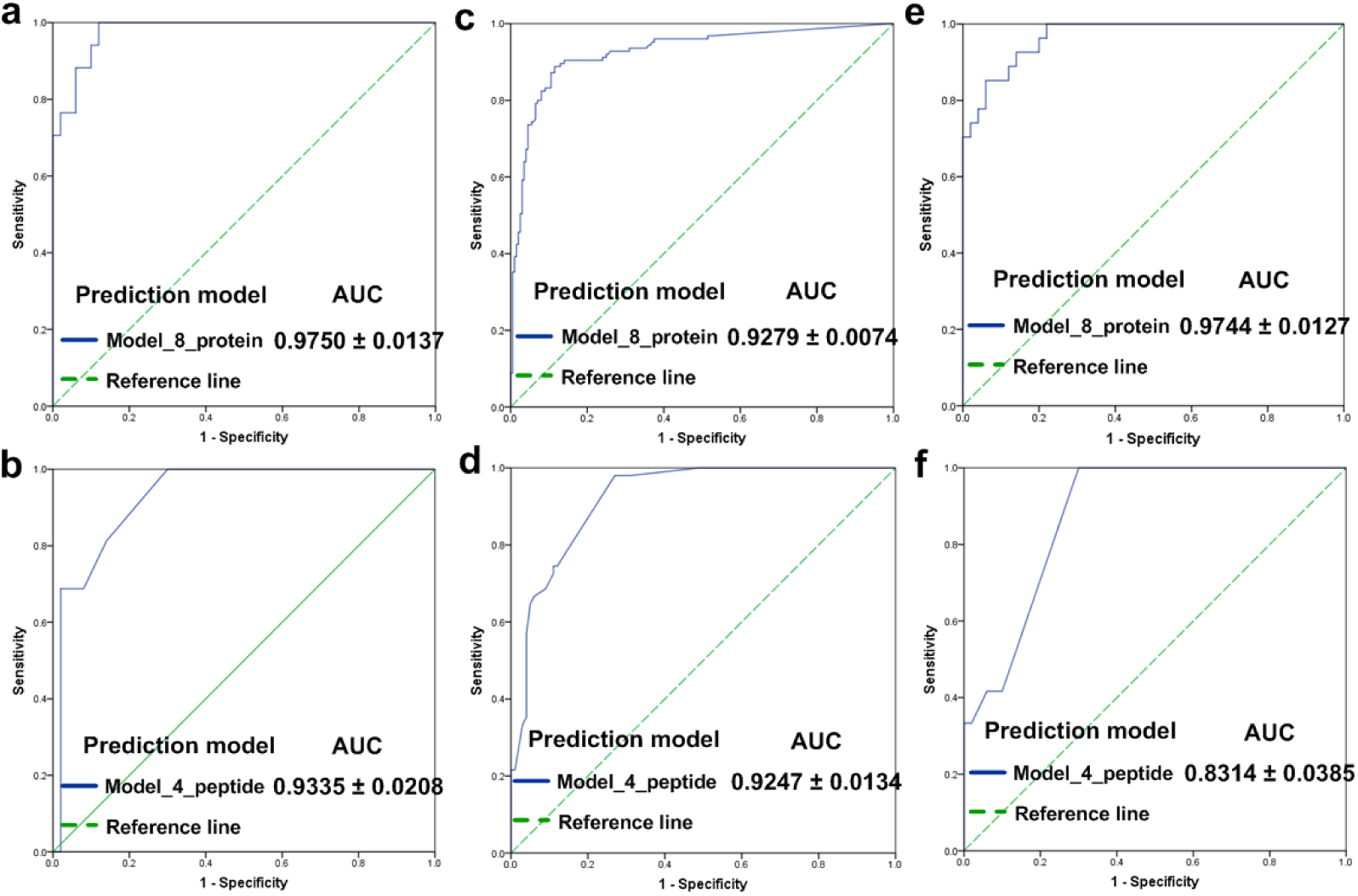
ROC curves and their corresponding AUCs of “Model_8_protein” and “Model_4_peptide” based on the independent test set1 (**a, b**), set2 (**c, d**) and set3 (**e, f**) (see Materials and Methods for more details).

## Discussion

In the field of drug discovery, systematic exploration of target space in the human genome always draws high attention (e.g. 14,15,19,21,27,57). However, owing to development history of the pharmaceutical industry, previous studies focused on either traditional small-molecule drugs’ targets, or all targets regardless of different types of drugs. Although some researchers have realized that targets of different types of drugs should be treated differently (23∼25), so far no one has systematically identified the similarities and differences between targets of protein, peptide and small-molecule drugs, let alone established target prediction models specially for protein/peptide drugs. Here we systematically analyzed and compared known targets of protein and peptide drugs, and presented distinctive characteristics of targets of protein and peptide drugs in multiple aspects and pointed out the difference between targets of protein, peptide and small-molecule drugs. Because of this difference, previous target prediction models based on analyses of small-molecule drugs’ targets or all targets regardless of drug types are not proper for or even will mislead the target selection of protein/peptide drugs. Therefore then based on these distinctive features, we constructed the first effective genome-wide target estimation models specially for protein and peptide drugs. This work not only contributes to the understanding of distinctive cellular roles of targets and the molecular mechanism of action of protein/peptide drugs from an overall perspective, but also more importantly provides the algorithm to give rational, quantitative advice for further target selection, helping improve the efficiency and success rate of target selection and validation, and ultimately contribute to novel target discovery for protein/peptide drugs.

## Acknowledgements

We would like to thank Jiannan Feng, He Xiao, Yuanfeng Li, Congwen Wei, Di Liu, Dan Wang, Liang Lu and Lihong Diao for fruitful discussion.

