## supplementary information for "Exploration of target spaces in the human genome for protein and peptide drugs"

**for**

**Table of Contents**


Supplementary Materials and Methods--------------------------------------------------------3

Supplementary Figure 1----------------------------------------------------------------------8

Supplementary Figure 2----------------------------------------------------------------------9

Supplementary Figure 3---------------------------------------------------------------------11

Supplementary Figure 4---------------------------------------------------------------------14

Supplementary Figure 5---------------------------------------------------------------------16

Supplementary Table 4----------------------------------------------------------------------17

Supplementary Table 5----------------------------------------------------------------------18

Supplementary Table 6----------------------------------------------------------------------19

Supplementary Table 7----------------------------------------------------------------------20

Supplementary Table 8----------------------------------------------------------------------21

Supplementary Table 9----------------------------------------------------------------------22

Supplementary Table 10---------------------------------------------------------------------23

Supplementary Table 11---------------------------------------------------------------------24

Supplementary Table 12---------------------------------------------------------------------25

Supplementary Table 14---------------------------------------------------------------------26

Supplementary Table 15---------------------------------------------------------------------27

Supplementary References-------------------------------------------------------------------28

### Supplementary Materials and Methods

#### The tissue specificity score (TSPS)

The tissue specificity score (TSPS) was adopted to measure the degree of tissue specific expression of a gene (Ravasi *et al.*, 2010). TSPS is computed as:

$$\text{TSPS}=\sum_i f_i \log_2(f_i / p) \quad [1]$$

where $f_i$ is the gene expression level in tissue *i* divided by the sum of expression levels across all tissues, and $p$=1/$n$ ($n$ is the number of tissues). The larger TSPS value is, the more tissue-specific the expression of the gene is.

#### Degree and Betweenness centrality

The degree of a node in the network is equal to the number of its interaction partners. Betweenness centrality is computed as:

$$B(n)=\frac{2\times\sum_{s\neq n\neq t}\frac{\sigma_{st}(n)}{\sigma_{st}}}{(N-1)(N-2)} \quad [2]$$

where $\sigma_{st}$ represents the number of the shortest paths between node *s* and *t*, $\sigma_{st}(n)$ is the number of those paths that go through node *n*, *N* is the total number of nodes in the network, and thus ($N$-1)($N$-2)/2 is the number of all possible node pairs (excluding node *n*) in the network which is also the maximum possible value of $\sum_{s\neq n\neq t}\frac{\sigma_{st}(n)}{\sigma_{st}}$ (Liu *et al*., 2013).

#### Data sources and methods related to Fig. 1d in the main document

To analyze the drug-disease relationship (Fig. 1d in the main document), we used high-confident drug-disease association data provided by Guney *et al.* (2016), which integrated data from MEDication Indication high precision subset (MEDI-HPS) (Wei *et al*., 2013), Kyoto Encyclopedia of Genes and Genomes (KEGG) (Kanehisa *et al*.,

2017) and National Drug File - Resource Terminology (NDF-RT) (https://www.nlm.nih.gov/research/umls/sourcereleasedocs/current/NDFRT/), and were strictly controlled on the quality. The approved drug-therapeutic target interaction data were from DrugBank (downloaded on 07/26/2015) as stated in the main document. As for the known gene-disease associations, we used Menche *et al.*'s scheme (2015) to combine data from UniProtKB/Swiss-Prot (gene-OMIM disease associations, downloaded on 3/23/2016) and Phenotype-Genotype Integrator (PheGenI) (downloaded on 10/21/2016) (Ramos *et al*., 2014).

**Minimum redundancy maximum relevance (mRMR) feature selection**

We used mRMR method for features selection. mRMR ranks features based on both their relevance to the classification variable and the redundancy between each other (Liu *et al*., 2013; Ding and Peng, 2005). Both the relevance and redundancy are quantified by mutual information (MI). MI denoted by $I$ between two discrete random variables $X$ and $Y$ is computed as:

$$I(X,Y)=\sum_{y\in Y}\sum_{x\in X}p(x,y)\log\left(\frac{p(x,y)}{p(x)p(y)}\right) \qquad [3]$$

where $p(x, y)$ is the joint probabilistic distribution, and $p(x)$ and $p(y)$ are respectively marginal probability distribution functions. Mutual information measures the mutual dependence between two random variables, that is, it quantifies the amount of information obtained about one random variable through the other random variable.

Suppose $S$ and $T$ are separately the already-selected and to-be-selected feature sets and $|S|$ and $|T|$ are the number of features in corresponding sets. mRMR firstly selects the feature most relevant to the classification variable into $S$, and then moves the remaining features from $T$ into $S$ one by one, requiring that each time the selected feature $f_j$ optimizes:

$$\max_{f_j \in T}\left( I(f_j, c) - \frac{1}{|S|}\sum_{f_i \in S} I(f_j, f_i) \right) \quad [4]$$

where $I(f_j, c)$ is the relevance of the candidate feature $f_j$ to the classification variable $c$ and $\frac{1}{|S|}\sum_{f_i \in S} I(f_j, f_i)$ is its redundancy with features already in $S$.

The GSN set was repeatedly constructed 100 times as stated above, and thus the mRMR feature ranking was implemented based on mean mutual information (MI) of 100 times between pairs of features and that between every feature and the classification variable.

The mRMR program was obtained from http://home.penglab.com/proj/mRMR/.

**Naïve Bayes classifier**

According to the Bayes rule, the posterior odds $O_{post}$ of a protein to be a target can be computed as the product of its prior odds $O_{piror}$ and the likelihood ratio (LR) represented by equation [5~7], where $P(positive)$ and $P(negative)$ are respectively the probability that a protein is and is not a drug target, and $P(positive \mid f_1 \ldots f_n)$ and $P(negative \mid f_1 \ldots f_n)$ are the corresponding probability after considering $n$ biological evidence types (i.e. features).

$$O_{post} = O_{prior} \times \mathrm{LR}(f_1 \ldots f_n) \quad [5]$$

$$O_{piror} = \frac{P(positive)}{P(negative)} \quad [6]$$

$$O_{post} = \frac{P(positive \mid f_1 \ldots f_n)}{P(negative \mid f_1 \ldots f_n)} \quad [7]$$

The LR of biological evidence $f$ is referred to as the ratio of the probability of feature $f$ observed in the GSP set to that in the GSN set:

$$\mathrm{LR}(f) = \frac{P(f \mid positive)}{P(f \mid negative)} = \frac{TP_f / T}{FP_f / F} \quad [8]$$

where $T$ and $F$ are the number of true and false drug targets in the golden standard dataset, and $TP_f$ and $FP_f$ are the number of those satisfying evidence $f$. LR can reflect the prediction ability of biological evidence $f$. When LR is larger than 1, $O_{post}$ is larger than $O_{piror}$, which means that the feature has the ability to discriminate true drug targets from the false ones. When the integrated $n$ features are independent, the naïve Bayes rule can be adopted, where the combined_LR can be calculated simply as the product of LRs of individual features (equation [9]).

$$\mathrm{LR}(f_1 \ldots f_n) = \prod_{i=1}^{i=n} \mathrm{LR}(f_i) \quad [9]$$

Actually the prior odds $O_{piror}$ is a constant for different proteins, and thus $O_{post}$ is proportional to the LR. In practice we directly used LR as the prediction score reflecting the probability that a protein is a drug target (Li *et al.*, 2008; Rhodes *et al.*, 2005).

Naïve Bayes classifier was implemented by Perl scripts written in-house.

**Performance assessment**

We used Receiver Operating Characteristic (ROC) curve to measure the performance of the prediction model, based on 10-fold cross-validation and independent test sets.

ROC curve shows the sensitivity and specificity of the prediction model against varying score cutoffs, whose *x*-axis is 1-specificity and *y*-axis sensitivity. Sensitivity ($TP/T$) and specificity ($1-FP/F$) are respectively correctly classified fraction in the positive ($T$) and negative test set ($F$), and $TP$ (true positive) and $FP$ (false positive) are the predicted positives in $T$ and $F$. A good prediction model has a ROC curve climbing rapidly toward the top left corner of the graph, which can be measured by the area under the curve (AUC). A perfect classifier has an AUC of 1.0, while

AUC=0.5 means that the prediction model is non-informative. Larger AUC indicates stronger prediction power (Liu *et al.*, 2013; Li *et al.*, 2008; Liu *et al.*, 2015). Here ROC curves were plotted by SPSS software (SPSS, Inc., 1999).

In 10-fold cross-validation, the golden standard dataset was averagely and randomly divided into 10 subsets. 9 ones were merged to be used as the training set to compute LRs of features, and the remaining one as the test set to count the number of *TP* and *FP* at different $LR_{cutoff}$. If the LR of a protein given by the prediction model exceeded $LR_{cutoff}$, it was viewed as the predicted positive. This process was repeated 10 times, and the numbers of TPs and FPs against varying $LR_{cutoff}$ of 10 test sets were summed to ultimately draw the ROC curve (Liu *et al.*, 2013; Li *et al.*, 2008; Liu *et al.*, 2015).

As for the assessment by the independent test set, the whole golden standard set was used as the training set, and the independent test set for plotting the ROC curve.

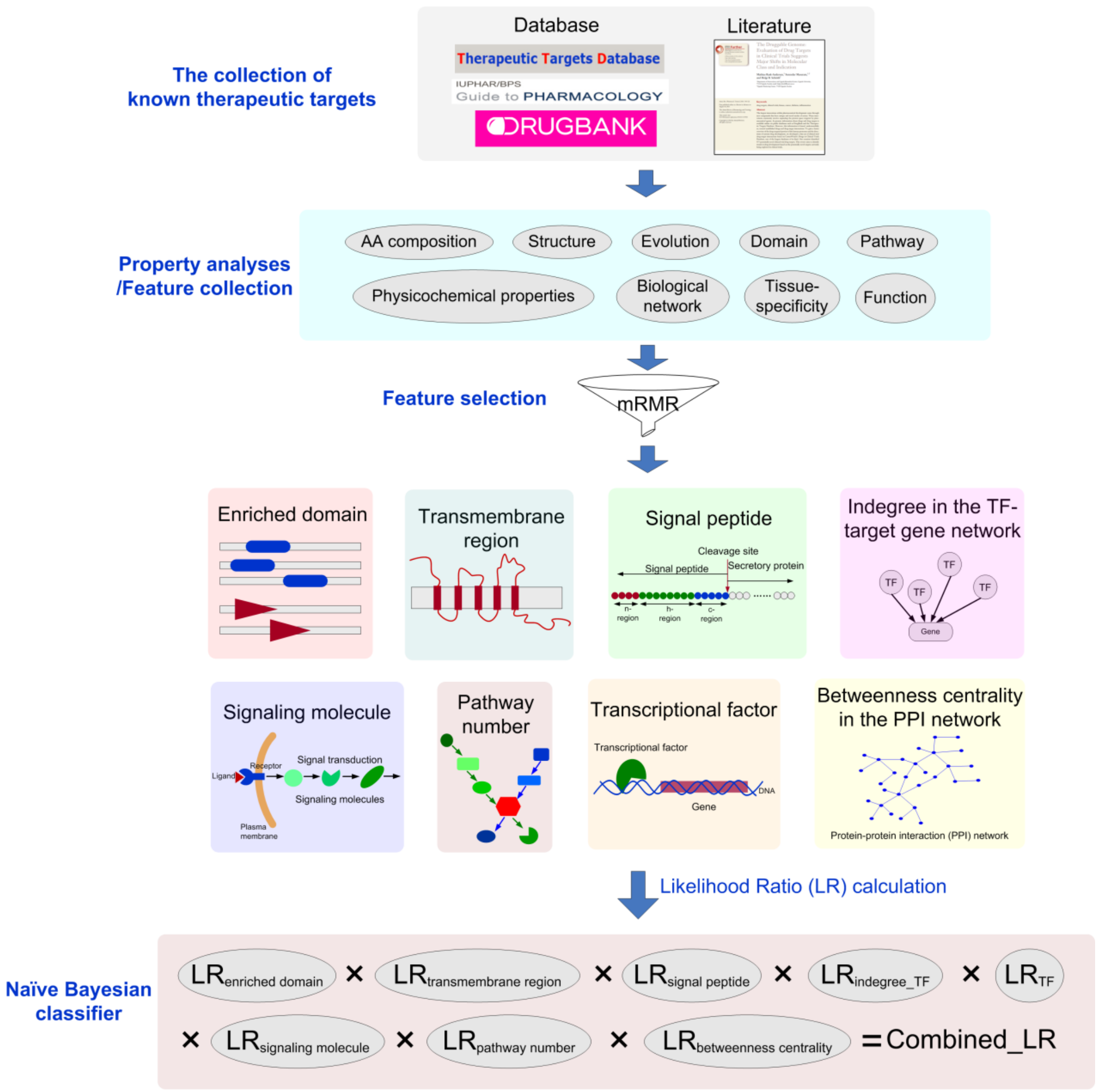


**Supplementary Figure 1** Workflow to explore target space for protein drugs. Workflow for peptide drugs is similar.

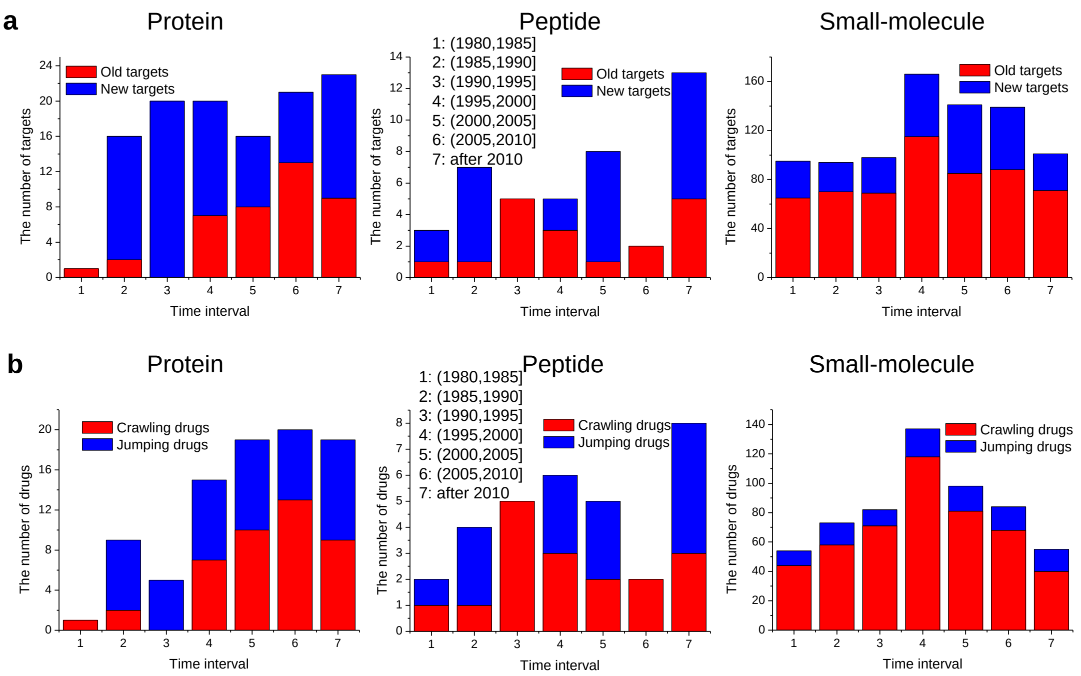


**Supplementary Figure 2** The composition of new drugs and their targets approved each five years based on approved drugs' therapeutic target data from DrugBank (downloaded on 07/26/2015) (see Materials and Methods in the main document). **a** The composition of targets of new drugs approved each five years. "Old targets" are referred to as those that have been used as targets of previously approved drugs, while "new targets" not. Each five years is a bin, and the year scope of each bin is described on the graph. The average proportion of "new targets" for protein drugs is significantly larger than that for small-molecule drugs (57.35%>31.93%, rank sum test, P-value=0.0380), while there is no statistically significant difference between peptide and small-molecule drugs (48.77%>31.93%, P-value=0.2090). **b** The composition of new drugs approved each five years. "Jumping drugs" are those with totally new target sets, whereas "crawling drugs" with at least one "old target" (Yildirim *et al.*, 2007). The average percentage of "jumping drugs" for protein drugs is significantly larger than that for small-molecule drugs (52.30%>18.57%, rank sum test, P-value=0.0260), while there is no statistically significant difference between peptide and small-molecule drugs (42.50%>18.57%, P-value=0.2090).

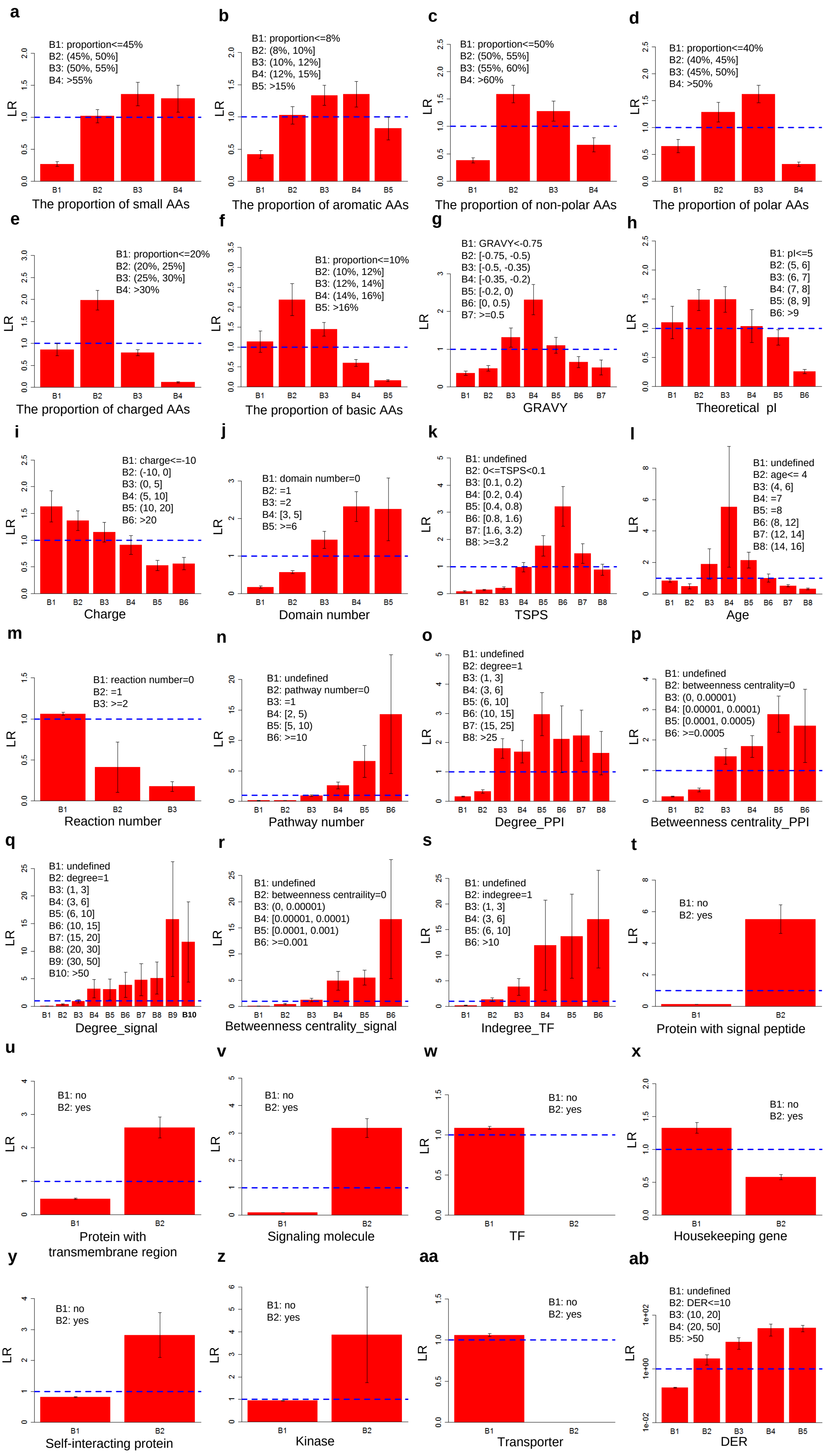
a
B1: proportion<=45%
B2: (45%, 50%]
B3: (50%, 55%]
B4: >55%
LR
The proportion of small AAs
b
B1: proportion<=8%
B2: (8%, 10%]
B3: (10%, 12%]
B4: (12%, 15%]
B5: >15%
The proportion of aromatic AAs
c
B1: proportion<=50%
B2: (50%, 55%]
B3: (55%, 60%]
B4: >60%
The proportion of non-polar AAs
d
B1: proportion<=40%
B2: (40%, 45%]
B3: (45%, 50%]
B4: >50%
The proportion of polar AAs
e
B1: proportion<=20%
B2: (20%, 25%]
B3: (25%, 30%]
B4: >30%
The proportion of charged AAs
f
B1: proportion<=10%
B2: (10%, 12%]
B3: (12%, 14%]
B4: (14%, 16%]
B5: >16%
The proportion of basic AAs
g
B1: GRAVY<-0.75
B2: [-0.75, -0.5)
B3: [-0.5, -0.35)
B4: [-0.35, -0.2)
B5: [-0.2, 0)
B6: [0, 0.5)
B7: >=0.5
GRAVY
h
B1: pI<=5
B2: (5, 6]
B3: (6, 7]
B4: (7, 8]
B5: (8, 9]
B6: >9
Theoretical pI
i
B1: charge<=-10
B2: (-10, 0]
B3: (0, 5]
B4: (5, 10]
B5: (10, 20]
B6: >20
Charge
j
B1: domain number=0
B2: =1
B3: =2
B4: [3, 5]
B5: >=6
Domain number
k
B1: undefined
B2: 0<=TSPS<0.1
B3: [0.1, 0.2)
B4: [0.2, 0.4)
B5: [0.4, 0.8)
B6: [0.8, 1.6)
B7: [1.6, 3.2)
B8: >=3.2
TSPS
l
B1: undefined
B2: age<= 4
B3: (4, 6]
B4: =7
B5: =8
B6: (8, 12]
B7: (12, 14]
B8: (14, 16]
Age
m
B1: reaction number=0
B2: =1
B3: >=2
Reaction number
n
B1: undefined
B2: pathway number=0
B3: =1
B4: [2, 5)
B5: [5, 10)
B6: >=10
Pathway number
o
B1: undefined
B2: degree=1
B3: (1, 3]
B4: (3, 6]
B5: (6, 10]
B6: (10, 15]
B7: (15, 25]
B8: >25
Degree_PPI
p
B1: undefined
B2: betweenness centrality=0
B3: (0, 0.00001)
B4: [0.00001, 0.0001)
B5: [0.0001, 0.0005)
B6: >=0.0005
Betweenness centrality_PPI
q
B1: undefined
B2: degree=1
B3: (1, 3]
B4: (3, 6]
B5: (6, 10]
B6: (10, 15]
B7: (15, 20]
B8: (20, 30]
B9: (30, 50]
B10: >50
Degree_signal
r
B1: undefined
B2: betweenness centraility=0
B3: (0, 0.00001)
B4: [0.00001, 0.0001)
B5: [0.0001, 0.001)
B6: >=0.001
Betweenness centrality_signal
s
B1: undefined
B2: indegree=1
B3: (1, 3]
B4: (3, 6]
B5: (6, 10]
B6: >10
Indegree_TF
t
B1: no
B2: yes
Protein with signal peptide
u
B1: no
B2: yes
Protein with transmembrane region
v
B1: no
B2: yes
Signaling molecule
w
B1: no
B2: yes
TF
x
B1: no
B2: yes
Housekeeping gene
y
B1: no
B2: yes
Self-interacting protein
z
B1: no
B2: yes
Kinase
aa
B1: no
B2: yes
Transporter
ab
B1: undefined
B2: DER<=10
B3: (10, 20]
B4: (20, 50]
B5: >50
DER

**Supplementary Figure 3** Likelihood Ratios (LRs) of various features used for target prediction for protein drugs. LR of feature *f* is defined as the ratio of the proportion of proteins satisfying feature *f* in the golden standard positive (GSP) set to that in the golden standard negative (GSN) set. Generally LR>1 means that the feature has the prediction ability, which is marked by the blue dashed line on the graphs. The GSN set was repeatedly constructed 100 times (see Materials and Methods in the main document), and thus LRs presented in these figures are mean and standard derivation (denoted by error bar) of 100 times. **a~f** The proportion of small, aromatic, non-polar, polar, charged and basic amino acids (AAs) in the protein sequence. **g** The grand average of hydropathy (GRAVY) (see Materials and Methods in the main document). **h** Theoretical isoelectric point (pI). **i** Charge. **j** Domain number. Here the number of domains (including repeats) rather than that of domain types of a protein was counted. **k** The tissue specificity score (TSPS) (see Supplementary Materials and Methods in this document). The bin of B1 includes proteins without computed TSPS because of the lack of corresponding gene expression profile data. **l** Original age. Here proteins were grouped into 16 age classes, from the oldest “Cellular organisms” class of age 16 composed of proteins that originated from the common ancestor of three domains of tree of life (Eukiaryota, Bacteria and Archaea) to the youngest “Homo sapiens” class of age 1 whose proteins are only found in human. Proteins of age from 2 to 15 are respectively those that originated from the ancestor of Homininae, Euarchontoglires, Eutheria, Theria, Mammalia, Amniota, Euteleostomi, Chordata, Coelomata, Bilateria, Eumetazoa, Metazoa, Fungi/Metazoa group, Eukaryota (Please refer to our previous work (Liu *et al.*, 2011) for more details). **m, n** Reaction number and pathway number a protein participates in. **o, p** Degree and betweenness centrality in the protein-protein interaction (PPI) network. **q, r** Degree and betweenness centrality in the signal transduction network. **s** Indegree in the transcriptional regulation network. Indegree of a gene is the number of TFs regulating the (target) gene. **t** Protein with signal peptide. **u** Protein with transmembrane region. **v** Signaling molecule. **w** Transcriptional factor (TF). **x** Housekeeping gene. **y** Self-interacting protein. **z** Kinase. **aa** Transporter. **ab** Domain enrichment ratio (DER). If a protein has multiple enriched domains, the largest DER is used.

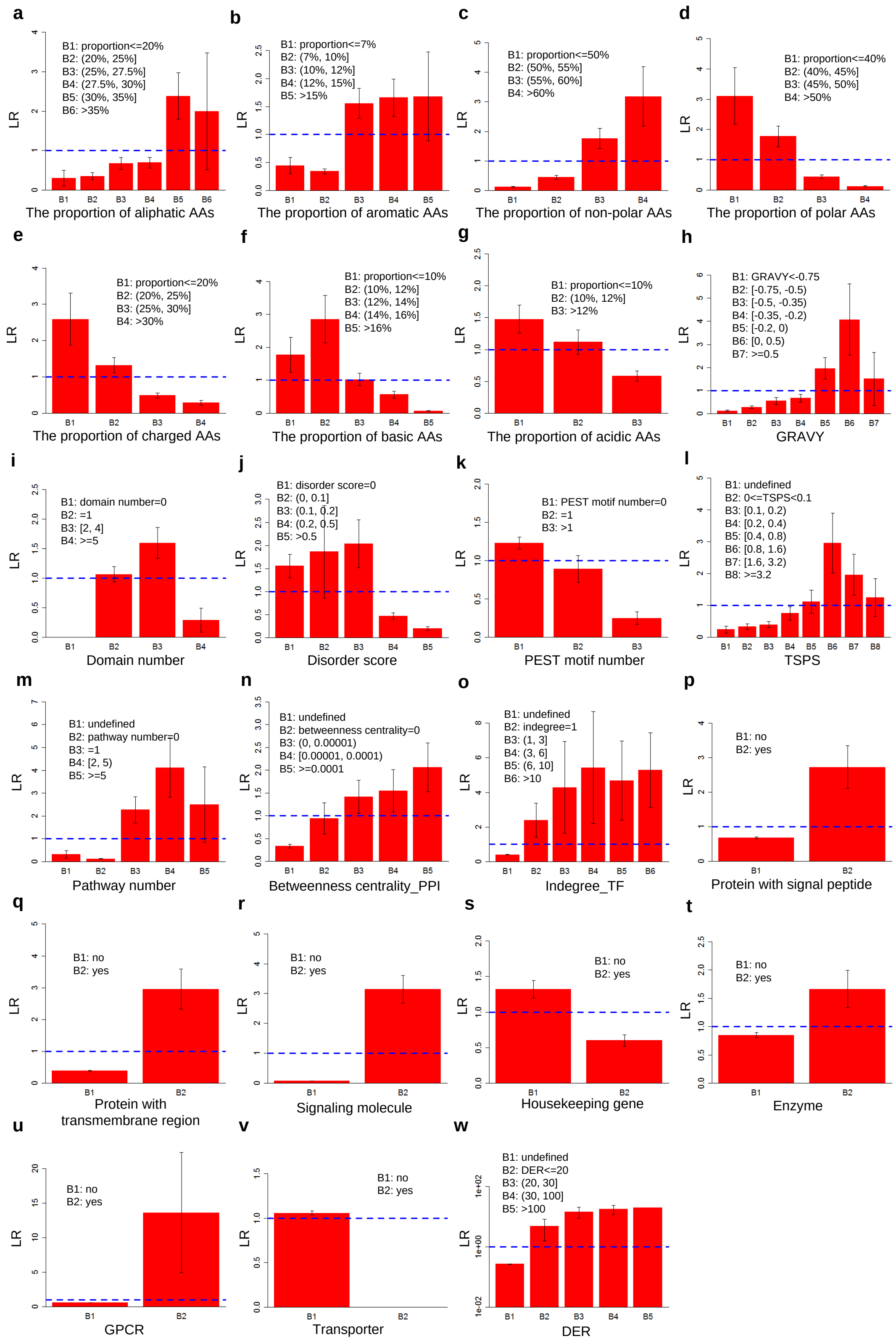


**Supplementary Figure 4** Likelihood Ratios (LRs) of various features used for target prediction for peptide drugs. Generally LR>1 means that the feature has the prediction ability, which is marked by the blue dashed line on the graphs. The GSN set was repeatedly constructed 100 times, and thus LRs presented in these figures are mean and standard derivation (denoted by error bar) of 100 times. **a~g** The proportion of aliphatic, aromatic, non-polar, polar, charged, basic and acidic amino acids (AAs) in the protein sequence. **h** The grand average of hydropathy (GRAVY). **i** Domain number. Here the number of domains (including repeats) rather than that of domain types of a protein was counted. **j** Disorder score. **k** PEST motif number. **l** The tissue specificity score (TSPS). **m** Pathway number a protein participates in. **n** Betweenness centrality in the PPI network. **o** Indegree in the transcriptional regulation network. Indegree of a gene is the number of TFs regulating the (target) gene. **p** Protein with signal peptide. **q** Protein with transmembrane region. **r** Signaling molecule. **s** Housekeeping gene. **t** Enzyme. **u** G protein-coupled receptor (GPCR). **v** Transporter. **w** Domain enrichment ratio (DER).

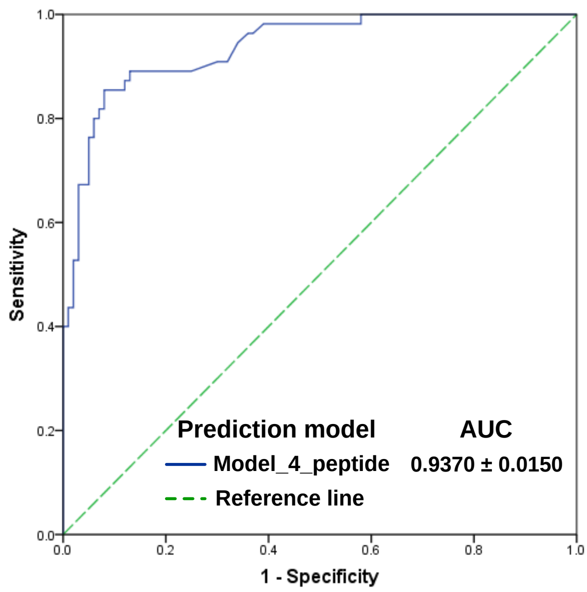


**Supplementary Figure 5** ROC curve and corresponding AUC of “Model_4_peptide” for target prediction for peptide drugs, integrating four features including DER, “signal peptide”, “transmembrane region” and “signaling molecule”, based on 10-fold cross-validation. The GSN set was repeatedly constructed 100 times, and thus mean ± SD of AUCs of 100 times is presented in the figure. The ROC curve in the figure was drawn based on GSP and a random GSN.

**Supplementary Table 4** The difference of quantitative properties between protein and small-molecule drugs' targets

| Property | Mean value (mean rank) | | P-value (rank sum test, one-sided) [a] |
|---|---|---|---|
| | Protein drugs' targets | Small-molecule drugs' targets | |
| Tiny (%) | 30.2151(435) | 28.8491 (373) | **0.0017** |
| Small (%) | 52.0710 (464) | 49.7220 (367) | **<0.0001** |
| Aliphatic (%) | 27.1799 (285) | 29.6454 (404) | **<0.0001** |
| Aromatic (%) | 10.8972 (314) | 11.9127 (398) | **<0.0001** |
| Non-polar (%) | 54.4961 (311) | 56.1784 (398) | **<0.0001** |
| Polar (%) | 45.5039 (456) | 43.8216 (369) | **<0.0001** |
| Charged (%) | 23.1265 (360) | 23.6275 (388) | 0.0854 |
| Basic (%) | 12.2549 (325) | 12.9508 (396) | **0.0004** |
| Acidic (%) | 10.8716 (392) | 10.6766 (382) | 0.3156 |
| GRAVY | -0.2802 (303) | -0.1469 (400) | **<0.0001** |
| Theoretical pI | 6.5670 (304) | 7.2178 (400) | **<0.0001** |
| Charge | -1.5644 (300) | 4.4968 (401) | **<0.0001** |
| Domain number | 3.5076 (432) | 2.3991 (373) | **0.0019** |
| Disorder score | 0.1984 (437) | 0.1575 (372) | **0.0010** |
| PEST motif number | 0.6591 (399) | 0.6215 (380) | 0.1509 |
| TSPS | 1.3331 (390) | 1.4096 (378) | 0.2878 |
| Age | 8.9126 (192) | 11.4509 (330) | **<0.0001** |
| Evolutionary rate | 4.5012 (369) | 2.0671 (270) | **<0.0001** |
| $C_{ratio}$ | 32.0566 (394) | 16.6069 (337) | **0.0026** |
| Pathway number | 6.7252 (441) | 5.8468 (370) | **0.0004** |
| Reaction number | 0.0303 (312) | 1.0599 (398) | **<0.0001** |
| Degree_PPI | 12.5484 (326) | 20.2363 (303) | 0.1030 |
| Betweenness centrality_PPI | 0.0003 (333) | 0.0009 (302) | **0.0375** |
| Degree_signal | 44.4797 (345) | 34.6157 (268) | **<0.0001** |
| Betweenness centrality_signal | 0.0013 (332) | 0.0013 (271) | **0.0002** |
| Indegree_TF | 8.3000 (298) | 6.5815 (243) | **0.0002** |
| Outdegree_TF | 3.1667 (29) | 32.3944 (44) | **0.0175** |

[a] P-values smaller than 0.05 are represented in bold type.

**Supplementary Table 5** The difference of qualitative properties between protein and small-molecule drugs' targets

| Property | The fraction of proteins belonging to a certain protein class (%) | | P-value (Fisher's exact test, one-sided) [a] |
|---|---|---|---|
| | Protein drugs' targets | Small-molecule drugs' targets | |
| Protein with signal peptide | 87.88 | 28.39 | **<0.0001** |
| Protein with transmembrane region | 64.39 | 54.42 | **0.0219** |
| Signaling molecule | 93.18 | 70.19 | **<0.0001** |
| Transcriptional factor | 0.00 | 4.42 | **0.0045** |
| Housekeeping gene | 25.00 | 30.28 | 0.1335 |
| Self-interacting protein | 26.52 | 21.61 | 0.1332 |
| Enzyme | 21.97 | 44.01 | **<0.0001** |
| G protein-coupled receptor (GPCR) | 6.06 | 14.67 | **0.0036** |
| Ion channel | 0.00 | 19.40 | **<0.0001** |
| Nuclear hormone receptors (NHR) | 0.00 | 3.31 | **0.0178** |
| Kinase | 8.33 | 9.62 | 0.3936 |
| Transporter | 0.00 | 25.71 | **<0.0001** |

[a] P-values smaller than 0.05 are represented in bold type.

**Supplementary Table 6** The difference of quantitative properties between peptide drugs' targets and other proteins

| Property | Mean value (mean rank) | | P-value (rank sum test, one-sided) [a] |
|---|---|---|---|
| | Peptide drugs' targets | Other proteins | |
| Tiny (%) | 29.9837 (11059) | 29.6644 (10014) | 0.0973 |
| Small (%) | 50.6965 (11186) | 49.8939 (10014) | 0.0722 |
| Aliphatic (%) | 30.9087 (13980) | 27.6174 (10006) | **<0.0001** |
| Aromatic (%) | 11.6914 (12736) | 10.4953 (10010) | **0.0003** |
| Non-polar (%) | 58.3613 (15087) | 53.2691 (10003) | **<0.0001** |
| Polar (%) | 41.6387 (4998) | 46.7305 (10031) | **<0.0001** |
| Charged (%) | 21.9288 (5925) | 25.4472 (10028) | **<0.0001** |
| Basic (%) | 11.8091 (5796) | 14.1326 (10029) | **<0.0001** |
| Acidic (%) | 10.1198 (7258) | 11.3145 (10025) | **0.0002** |
| GRAVY | -0.0235 (15152) | -0.3488 (10003) | **<0.0001** |
| Theoretical pI | 7.4680 (10608) | 7.3276 (10013) | 0.2343 |
| Charge | 4.5455 (10275) | 4.7546 (10016) | 0.3848 |
| Domain number | 1.7273 (11280) | 1.9429 (10014) | **0.0455** |
| Disorder score | 0.1180 (6600) | 0.2659 (10025) | **<0.0001** |
| PEST motif number | 0.2364 (8332) | 0.7310 (10022) | **0.0047** |
| TSPS | 1.4986 (11722) | 1.0878 (9113) | **0.0002** |
| Age | 10.2561 (7039) | 10.5679 (7424) | 0.2722 |
| Evolutionary rate | 0.2776 (6557) | 3.7016 (6774) | 0.3490 |
| $C_{ratio}$ | 27.4976 (7417) | 31.7672 (8003) | 0.1800 |
| Pathway number | 3.1852 (14967) | 1.3012 (9319) | **<0.0001** |
| Reaction number | 0.2909 (10552) | 0.4310 (10016) | 0.0739 |
| Degree_PPI | 8.6875 (6680) | 10.7513 (6176) | 0.1758 |
| Betweenness centrality_PPI | 0.0002 (7109) | 0.0002 (6174) | **0.0383** |
| Degree_signal | 23.6538 (3119) | 18.1905 (2997) | 0.3227 |
| Betweenness centrality_signal | 0.0003 (3279) | 0.0004 (2995) | 0.1286 |
| Indegree_TF | 4.1892 (2441) | 3.6237 (2047) | **0.0204** |
| Outdegree_TF | 14.7500 (516) | 15.5899 (497) | 0.4498 |

[a] P-values smaller than 0.05 are represented in bold type.

**Supplementary Table 7** The difference of qualitative properties between peptide drugs' targets and other proteins

| Property | The fraction of proteins belonging to a certain protein class (%) | | P-value (Fisher's exact test, one-sided) [a] |
|---|---|---|---|
| | Peptide drugs' targets | Other proteins | |
| Protein with signal peptide | 43.64 | 16.94 | **<0.0001** |
| Protein with transmembrane region | 70.91 | 25.48 | **<0.0001** |
| Signaling molecule | 94.55 | 29.76 | **<0.0001** |
| Transcriptional factor | 1.82 | 8.23 | 0.0528 |
| Housekeeping gene | 25.45 | 43.58 | **0.0043** |
| Self-interacting protein | 14.55 | 9.78 | 0.1662 |
| Enzyme | 32.73 | 20.17 | **0.0199** |
| G protein-coupled receptor (GPCR) | 41.82 | 3.85 | **<0.0001** |
| Ion channel | 0.00 | 1.62 | 0.4074 |
| Nuclear hormone receptors (NHR) | 1.82 | 0.23 | 0.1213 |
| Kinase | 0.00 | 2.49 | 0.2500 |
| Transporter | 0.00 | 5.68 | **0.0404** |

[a] P-values smaller than 0.05 are represented in bold type.

**Supplementary Table 8** The difference of quantitative properties between peptide and small-molecule drugs' targets

| Property | Mean value (mean rank) | | P-value (rank sum test, one-sided) [a] |
|---|---|---|---|
| | Peptide drugs' targets | Small-molecule drugs' targets | |
| Tiny (%) | 29.9837 (409) | 28.8491 (339) | **0.0067** |
| Small (%) | 50.6965 (390) | 49.7220 (341) | **0.0419** |
| Aliphatic (%) | 30.9087 (412) | 29.6454 (339) | **0.0046** |
| Aromatic (%) | 11.6914 (344) | 11.9127 (345) | 0.4808 |
| Non-polar (%) | 58.3613 (449) | 56.1784 (336) | **<0.0001** |
| Polar (%) | 41.6387 (241) | 43.8216 (354) | **<0.0001** |
| Charged (%) | 21.9288 (254) | 23.6275 (353) | **0.0002** |
| Basic (%) | 11.8091 (250) | 12.9508 (353) | **0.0001** |
| Acidic (%) | 10.1198 (280) | 10.6766 (351) | **0.0056** |
| GRAVY | -0.0235 (427) | -0.1469 (338) | **0.0008** |
| Theoretical pI | 7.4680 (373) | 7.2178 (343) | 0.1378 |
| Charge | 4.5455 (348) | 4.4968 (345) | 0.4574 |
| Domain number | 1.7273 (284) | 2.3991 (350) | **0.0066** |
| Disorder score | 0.1180 (281) | 0.1575 (351) | **0.0062** |
| PEST motif number | 0.2364 (295) | 0.6215 (349) | **0.0100** |
| TSPS | 1.4986 (359) | 1.4096 (340) | 0.2463 |
| Age | 10.2561 (227) | 11.4509 (279) | **0.0200** |
| Evolutionary rate | 0.2776 (312) | 2.0671 (254) | **0.0083** |
| $C_{ratio}$ | 27.4976 (344) | 16.6069 (309) | 0.0979 |
| Pathway number | 3.1852 (291) | 5.8468 (349) | **0.0194** |
| Reaction number | 0.2909 (306) | 1.0599 (348) | **0.0209** |
| Degree_PPI | 8.6875 (262) | 20.2363 (271) | 0.3478 |
| Betweenness centrality_PPI | 0.0002 (267) | 0.0009 (270) | 0.4371 |
| Degree_signal | 23.6538 (214) | 34.6157 (253) | **0.0327** |
| Betweenness centrality_signal | 0.0003 (207) | 0.0013 (254) | **0.0120** |
| Indegree_TF | 4.1892 (206) | 6.5815 (220) | 0.2596 |
| Outdegree_TF | 14.7500 (31) | 32.3944 (38) | 0.2491 |

[a] P-values smaller than 0.05 are represented in bold type.

**Supplementary Table 9** The difference of qualitative properties between peptide and small-molecule drugs' targets

| Property | The fraction of proteins belonging to a certain protein class (%) | | P-value (Fisher's exact test, one-sided) [a] |
|---|---|---|---|
| | Peptide drugs' targets | Small-molecule drugs' targets | |
| Protein with signal peptide | 43.64 | 28.39 | **0.0150** |
| Protein with transmembrane region | 70.91 | 54.42 | **0.0122** |
| Signaling molecule | 94.55 | 70.19 | **<0.0001** |
| Transcriptional factor | 1.82 | 4.42 | 0.3087 |
| Housekeeping gene | 25.45 | 30.28 | 0.2793 |
| Self-interacting protein | 14.55 | 21.61 | 0.1434 |
| Enzyme | 32.73 | 44.01 | 0.0685 |
| G protein-coupled receptor (GPCR) | 41.82 | 14.67 | **<0.0001** |
| Ion channel | 0.00 | 19.40 | **<0.0001** |
| Nuclear hormone receptors (NHR) | 1.82 | 3.31 | 0.4630 |
| Kinase | 0.00 | 9.62 | **0.0049** |
| Transporter | 0.00 | 25.71 | **<0.0001** |

[a] P-values smaller than 0.05 are represented in bold type.

**Supplementary Table 10** The difference of quantitative properties between protein and peptide drugs' targets

| Property | Mean value (mean rank) | | P-value (rank sum test, one-sided) [a] |
|---|---|---|---|
| | Protein drugs' targets | Peptide drugs' targets | |
| Tiny (%) | 30.2151 (93) | 29.9837 (96) | 0.3428 |
| Small (%) | 52.0710 (97) | 50.6965 (87) | 0.1331 |
| Aliphatic (%) | 27.1799 (80) | 30.9087 (127) | **<0.0001** |
| Aromatic (%) | 10.8972 (88) | 11.6914 (108) | **0.0100** |
| Non-polar (%) | 54.4961 (79) | 58.3613 (129) | **<0.0001** |
| Polar (%) | 45.5039 (109) | 41.6387 (59) | **<0.0001** |
| Charged (%) | 23.1265 (101) | 21.9288 (77) | **0.0027** |
| Basic (%) | 12.2549 (98) | 11.8091 (86) | 0.0847 |
| Acidic (%) | 10.8716 (101) | 10.1198 (76) | **0.0021** |
| GRAVY | -0.2802 (80) | -0.0235 (128) | **<0.0001** |
| Theoretical pI | 6.5670 (85) | 7.4680 (115) | **0.0003** |
| Charge | -1.5644 (86) | 4.5455 (113) | **0.0011** |
| Domain number | 3.5076 (104) | 1.7273 (71) | **0.0001** |
| Disorder score | 0.1984 (104) | 0.1180 (70) | **<0.0001** |
| PEST motif number | 0.6591 (100) | 0.2364 (80) | **0.0035** |
| TSPS | 1.3331 (92) | 1.4986 (96) | 0.3199 |
| Age | 8.9126 (68) | 10.2561 (85) | **0.0117** |
| Evolutionary rate | 4.5012 (71) | 0.2776 (62) | 0.1053 |
| $C_{ratio}$ | 32.0566 (85) | 27.4976 (82) | 0.3418 |
| Pathway number | 6.7252 (103) | 3.1852 (69) | **<0.0001** |
| Reaction number | 0.0303 (91) | 0.2909 (101) | **0.0006** |
| Degree_PPI | 12.5484 (89) | 8.6875 (79) | 0.1230 |
| Betweenness centrality_PPI | 0.0003 (90) | 0.0002 (78) | 0.0949 |
| Degree_signal | 44.4797 (99) | 23.6538 (62) | **<0.0001** |
| Betweenness centrality_signal | 0.0013 (100) | 0.0003 (61) | **<0.0001** |
| Indegree_TF | 8.3000 (80) | 4.1892 (57) | **0.0027** |
| Outdegree_TF | 3.1667 (8) | 14.7500 (9) | 0.4020 |

[a] P-values smaller than 0.05 are represented in bold type.

**Supplementary Table 11** The difference of qualitative properties between protein and peptide drugs' targets

| Property | The fraction of proteins belonging to a certain protein class (%) | | P-value (Fisher's exact test, one-sided) [a] |
|---|---|---|---|
| | Protein drugs' targets | Peptide drugs' targets | |
| Protein with signal peptide | 87.88 | 43.64 | **<0.0001** |
| Protein with transmembrane region | 64.39 | 70.91 | 0.2468 |
| Signaling molecule | 93.18 | 94.55 | 0.5087 |
| Transcriptional factor | 0.00 | 1.82 | 0.2941 |
| Housekeeping gene | 25.00 | 25.45 | 0.5426 |
| Self-interacting protein | 26.52 | 14.55 | 0.0538 |
| Enzyme | 21.97 | 32.73 | 0.0882 |
| G protein-coupled receptor (GPCR) | 6.06 | 41.82 | **<0.0001** |
| Ion channel | 0.00 | 0.00 | - |
| Nuclear hormone receptors (NHR) | 0.00 | 1.82 | 0.2941 |
| Kinase | 8.33 | 0.00 | **0.0191** |
| Transporter | 0.00 | 0.00 | - |

[a] P-values smaller than 0.05 are represented in bold type.

**Supplementary Table 12** ROC AUCs of single-feature target prediction models for protein drugs based on 10-fold cross-validation

| Single-feature prediction model [a] | ROC AUC (mean ± SD) [a] |
|---|---|
| DER | 0.8595 ± 0.0076 |
| Betweenness centrality_signal | 0.8594 ± 0.0149 |
| Degree_signal | 0.8593 ± 0.0148 |
| Pathway number | 0.8324 ± 0.0132 |
| Indegree_TF | 0.8252 ± 0.0141 |
| Signal peptide | 0.8176 ± 0.0144 |
| Signaling molecule | 0.7807 ± 0.0170 |
| TSPS | 0.7142 ± 0.0237 |
| Betweenness centrality_PPI | 0.7060 ± 0.0248 |
| Degree_PPI | 0.6908 ± 0.0282 |
| Transmembrane region | 0.6579 ± 0.0169 |
| Basic | 0.6477 ± 0.0262 |
| Domain number | 0.6467 ± 0.0210 |
| Age | 0.6464 ± 0.0212 |
| Charged | 0.6299 ± 0.0230 |
| GRAVY | 0.6251 ± 0.0244 |
| Polar | 0.6116 ± 0.0278 |
| Non-polar | 0.6015 ± 0.0288 |
| pI | 0.5705 ± 0.0252 |
| Housekeeping gene | 0.5556 ± 0.0179 |
| Charge | 0.5543 ± 0.0249 |
| Self-interacting protein | 0.5521 ± 0.0125 |
| Small | 0.5511 ± 0.0299 |
| Aromatic | 0.5486 ± 0.0280 |
| TF | 0.5258 ± 0.0071 |
| Transporter | 0.5161 ± 0.0081 |
| Kinase | 0.5110 ± 0.0064 |
| Reaction number | 0.5091 ± 0.0084 |

[a] The GSN set was repeatedly constructed 100 times, and thus the presented ROC AUCs are mean ± standard derivation (SD) of results of 100 times. The single-feature prediction models are ranked based on the order of decreasing mean value.

**Supplementary Table 14** Features used for target prediction for protein drugs ranked by mRMR method based on the golden standard dataset

| Order | Feature [a] |
|---|---|
| 1 | DER [b] |
| 2 | Indegree_TF |
| 3 | Signal peptide |
| 4 | Signaling molecule |
| 5 | Transmembrane region |
| 6 | Pathway number |
| 7 | Transcriptional factor |
| 8 | Betweenness centrality_PPI |
| 9 | Betweenness centrality_signal |
| 10 | Transporter |
| 11 | Basic |
| 12 | TSPS |
| 13 | Reaction number |
| 14 | Domain number |
| 15 | Degree_signal |
| 16 | Self-interacting protein |
| 17 | Small |
| 18 | Housekeeping gene |
| 19 | Charged |
| 20 | pI |
| 21 | Age |
| 22 | Kinase |
| 23 | Aromatic |
| 24 | Polar |
| 25 | Degree_PPI |
| 26 | Charge |
| 27 | GRAVY |
| 28 | Non-polar |

[a] The GSN set was repeatedly constructed 100 times, and thus the mRMR feature ranking was implemented based on mean mutual information (MI) of 100 times between pairs of features and that between every feature and the classification variable (see Supplementary Materials and Methods in this document). [b] For DER, 10-fold cross-validation protocol was used. Nine tenths of the golden standard dataset was used to define the enriched domains together with corresponding DERs, and the remaining one as the test set (when a protein contains multiple enriched domains, the largest DER was used). The process was repeated ten times and finally ten test sets were merged to be used for mRMR analysis.

**Supplementary Table 15** Features used for target prediction for peptide drugs ranked by mRMR based on the golden standard dataset

| Order | Feature |
|---|---|
| 1 | DER |
| 2 | Signaling molecule |
| 3 | Transmembrane region |
| 4 | Signal peptide |
| 5 | Pathway number |
| 6 | Non-polar |
| 7 | Transporter |
| 8 | GPCR |
| 9 | Indegree_TF |
| 10 | Domain number |
| 11 | PEST motif number |
| 12 | Betweenness centrality_ PPI |
| 13 | Basic |
| 14 | Aromatic |
| 15 | Enzyme |
| 16 | Housekeeping gene |
| 17 | Disorder score |
| 18 | Acidic |
| 19 | Aliphatic |
| 20 | TSPS |
| 21 | Polar |
| 22 | Charged |
| 23 | GRAVY |
